# Early life Western diet-induced memory impairments and gut microbiome changes in female rats are long-lasting despite healthy dietary intervention

**DOI:** 10.1101/2021.04.05.438525

**Authors:** Linda Tsan, Shan Sun, Lana Bridi, Lekha S. Chirala, Emily E. Noble, Anthony A. Fodor, Scott E. Kanoski

**Affiliations:** Neuroscience Graduate Program, University of Southern California, Los Angeles, CA, USA; Department of Biological Sciences, Human and Evolutionary Biology Section, University of Southern California, Los Angeles, CA, USA; Department of Foods and Nutrition, University of Georgia, Athens, GA, USA; Department of Bioinformatics and Genomics, University of North Carolina at Charlotte, Charlotte, NC, USA

**Keywords:** Obesity, junk food, hippocampus, microbiota, high-fat diet, sugar

## Abstract

**Objective:** Consumption of a Western diet during adolescence results in hippocampus (HPC)-dependent memory impairments and gut microbiome dysbiosis. Whether these adverse outcomes are reversible in adulthood following intervention with a healthy diet is unknown. Here we assessed the short- and long-term effects of adolescent consumption of a Western diet enriched with either sugar alone, or sugar and fat on metabolic outcomes, HPC-dependent memory, and gut microbiota.

**Methods:** Adolescent female rats (PN 26) were fed a standard chow diet (CTL), a chow diet with access to 11% sugar solution (SUG), or a junk food cafeteria-style diet (CAF) containing a variety of fat- and/or sugar-enriched foods. During adulthood (PN 65+), metabolic outcomes, HPC-dependent memory, and gut microbial populations were evaluated both before and after a 5-week dietary intervention period where all groups were fed a diet of water standard chow.

**Results:** Prior to the dietary intervention both the CAF and SUG groups demonstrated impaired HPC-dependent memory, increased adiposity, and altered gut microbial populations relative to controls. However, impaired peripheral glucose regulation was only observed in the SUG group. The dietary intervention reversed the metabolic dysfunction in both the CAF and SUG groups, whereas HPC-dependent memory impairments were reversed in the SUG, but not the CAF group. The composition of the gut microbiota remained distinct from controls in both groups after dietary intervention.

**Conclusions:** While the metabolic impairments associated with adolescent cafeteria diet consumption are reversible in adulthood with dietary intervention, the HPC-dependent memory impairments and the gut microbiome dysbiosis persist.

## 1. Introduction

Globalization has brought technological advances in food processing, shelf-life, marketing, and distribution practices have increased the availability of low-cost palatable foods that have a high percentage of calories derived from saturated fat and sugar. The increased prevalence and accessibility of high fat, high sugar foods, herein referred to as a Western diet (WD), has directly impacted the diet quality of U.S. citizens, particularly children, as U.S. children on average exceed the recommended guidelines for consumption of sugars and fats before they reach school age (Wang et al., 2018). Furthermore, in U.S. children aged 2 to 19 years, ultra-processed foods rich in lipids and added sugars contribute to ∼65% of total energy intake (Neri et al., 2019). Indeed, the declining quality in the WD is likely contributing to the increasing rates of childhood obesity as consumption of ultra-processed foods in children is positively associated with body fat (Wang et al., 2015; Costa et al., 2018; Gallagher et al., 2020) and increased risk for metabolic syndrome (Ambrosini et al., 2010; Mager et al., 2013).

In addition to adverse metabolic outcomes, WD exposure during early life periods of development adversely impacts neurocognitive development (Francis and Stevenson, 2013; Noble and Kanoski, 2016; Reichelt, 2016; Tsan et al., 2021). For example, habitual added sugar consumption in children is associated with altered hippocampus (HPC) volume and HPC-cortical connectivity (Clark et al., 2020), whereas habitual saturated fatty acid intake is negatively associated with HPC-dependent relational memory (Baym et al., 2014). Rodent studies also reveal impaired HPC-dependent learning and memory associated with early life consumption of either a WD or excessive sugar consumption without elevated fat intake, even in cases where the WD (or sugar) consumption is not accompanied by metabolic dysfunction and/or obesity (Kendig et al., 2013; Hsu et al., 2015a; Ferreira et al., 2018; Noble et al., 2019). The overwhelming majority of these rodent model studies tested memory performance during adulthood while the animals were still being maintained on the WD or the high sugar diet. While memory impairments associated with WD consumption in male rats can be reversed with intervention, such as exercise (Noble et al., 2014), very little is understood about whether these early life WD-induced memory impairments can be reversed with healthy dietary intervention during adulthood, with one study finding that removing access to a sugar-sweetened beverage solution for 4 months in adulthood after prolonged access in adolescence with an otherwise standard rodent diet did not reverse sugar-sweetened beverage-induced HPC-dependent memory impairment in male rats (Noble et al., 2019). Additionally, few studies have specifically investigated whether dietary intervention reverses diet induced cognitive dysfunction in females.

Emerging evidence suggests that the gut microbiome influences cognitive function (Noble et al., 2017b; Provensi et al., 2019; Leigh et al., 2020; Darch et al., 2021; Noble et al., 2021) and that this relationship may be dependent on early life nutrition (Cerdó et al., 2019). For example, excessive early life sugar consumption in male rats yields HPC-dependent memory impairments (Hsu et al., 2015a), an outcome that was recently connected functionally to robust changes in the microbiome relative to controls (Noble et al., 2021). In another study, transplanting the microbiota of male mice that received an early life high fat diet to control mice conferred HPC-dependent learning and memory deficits, suggesting a possible functional connection between microbiota composition and HPC-dependent learning and memory (Yang et al., 2019). However, as is the case with WD-associated memory impairments, it is poorly understood whether microbiome outcomes linked to early life WD consumption can be reversed with healthy dietary intervention. In humans, after the first 3 years of life an estimated 60-70% of the microbiota composition remains relatively stable throughout life, yet ∼30-40% may be more susceptible to changes induced by diet or other factors (Kashtanova et al., 2016). The extent to which microbiome changes induced by dietary factors during early life developmental periods are long-lasting vs. reversible when the habitual diet changes during adulthood is unknown.

In the present study, we evaluate, neurocognitive (HPC-dependent memory, anxiety-like behavior), metabolic (glucose tolerance, body weight, adiposity ratio, caloric intake), and gut microbiome outcomes in female rats exposed to either control conditions (free access to water and standard low-fat rat chow) or one of two different WD models during the entire adolescent period of development: 1) a cafeteria-style junk food diet (CAF) group, with free choice access to water, a rodent high-fat diet, an 11% carbohydrate weight-by-volume (w/v) high fructose corn syrup (HFCS) solution, potato chips, and chocolate peanut butter cups, or 2) a group with access to water, standard low-fat rat chow, and the 11% w/v HFCS solution (SUG). In order to explore possible links between gut microbiota and memory outcomes, linear regression analyses were conducted to determine the relationship between bacterial taxa abundancies and HPC-dependent memory performance.

## 2. Materials and methods

### 2.1 Subjects and Diets

Adolescent (postnatal day [PN] 26) female Sprague Dawley rats (n=20, Envigo; 50-70g) were housed individually in a climate-controlled (22–24 °C) environment with a 12:12 reverse light/dark cycle and maintained on standard chow (Lab Diet 5001; PMI Nutrition International, Brentwood, MO; 29.8 % kcal from protein, 13.4% kcal from fat, 56.7% kcal from carbohydrate) and water unless otherwise stated. All experiments were approved by the Animal Care and Use Committee at the University of Southern California and performed in accordance with the National Research Council Guide for the Care and Use of Laboratory Animals.

### 2.2 Experiment 1 Design (Junk Food-Style Cafeteria Diet)

At PN 26, rats were randomized to one of two groups (matched for body weight) and received ad libitum access to either [1] standard chow and water (control [CTL] group, n=10), or [2] a cafeteria (CAF) diet, with free access to potato chips (Ruffles), peanut butter cups (Reese’s minis), 45% kcal high fat/sucrose chow (Research Diets, D12451)), a bottle of 11% weight/volume (w/v) high fructose corn syrup (HFCS)-55 solution, and water (CAF group, n=10). The concentration of sugar (11% w/v) was selected to model the amount of sugar present in sugar-sweetened beverages commonly consumed by humans, and is also based on our prior studies in male rats (Hsu et al., 2015b; Noble et al., 2019). Three food hoppers were used in each cage for precise measure of individual solid food types (in CAF group) or for consistency across experimental groups (in CTL group, all hoppers filled with standard chow). Papers were placed underneath the cages to enable the collection and recording of food spillage. Body weights and food intake were measured 3x/week throughout the study. At PN 61 (young adulthood), testing began for the Novel Object in Context (NOIC) task, which measures hippocampal-dependent episodic/contextual memory (Martínez et al., 2014). Anxiety-like behavior was assessed using a Zero Maze at PN 67. Fecal samples for analyses of gut microbiota were collected at PN 70 (Timepoint A, n=10/group). Body composition was measured using nuclear magnetic resonance (NMR) spectroscopy at PN 74 and an intraperitoneal glucose tolerance test (IPGTT) was conducted at PN 78. At PN 79, animals in CAF were switched to a standard chow diet with water access for the remainder of the study as a dietary intervention. Body composition was reevaluated at PN 115. A second cohort of female rats (n=31, 50-70g) were treated as described above in order to collect behavioral data following the dietary intervention. These rats were tested on NOIC after 5 weeks of dietary intervention (chow and water only) at PN 101 and Zero Maze at PN 108 (Figure 1B). Fecal samples for microbiota analyses were collected after the dietary intervention at PN 106 (Timepoint B, n=15 CTL group, n=16 CAF group). Caloric intake and body weight were recorded 3x/week and IPGTT conducted while still on the diet (PN 66).

**Figure 1:**
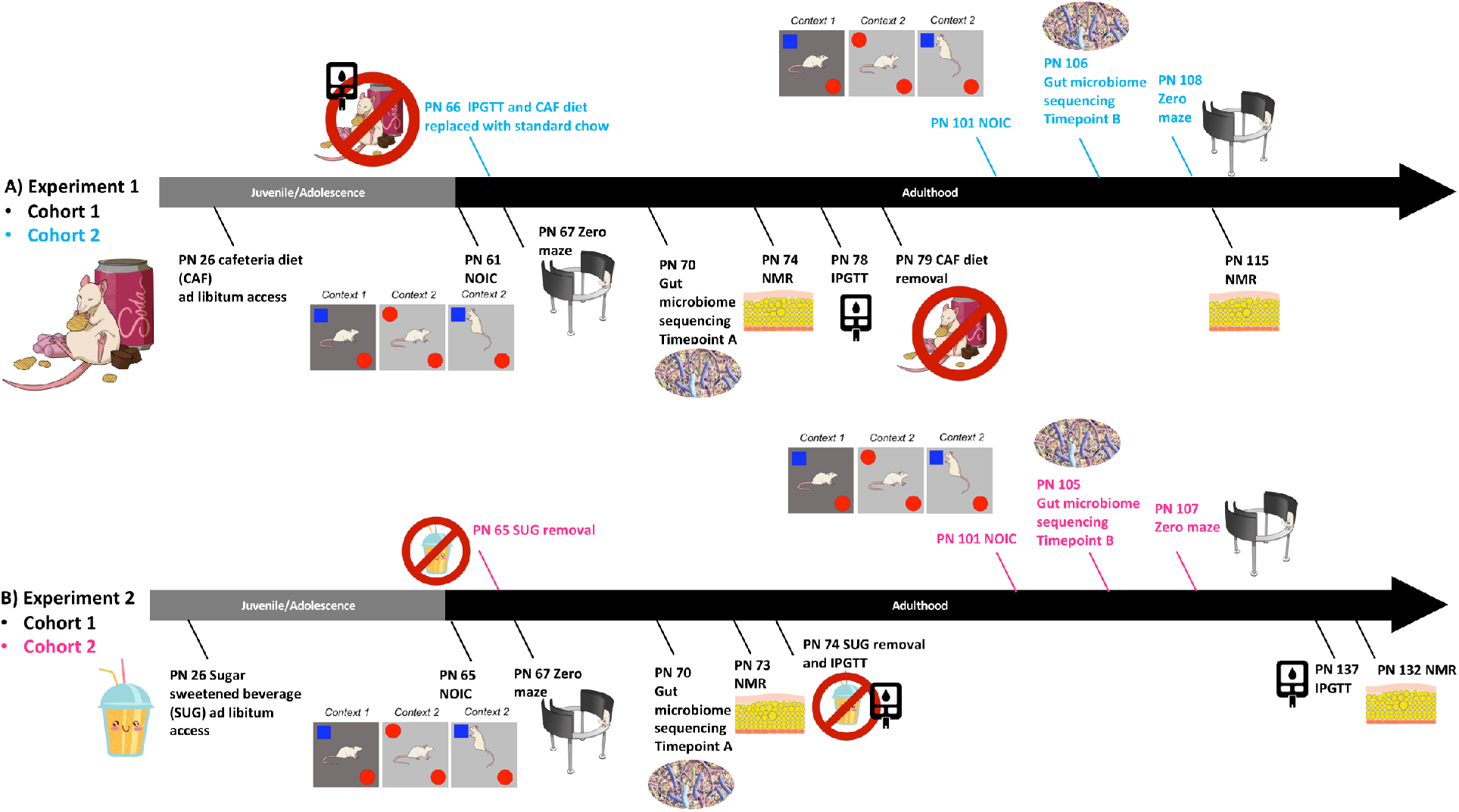
Timeline of experiments. During the adolescent period of development, rats were exposed to either a junk food-style cafeteria diet (A) or an otherwise healthy diet with access to a sugar-sweetened beverage (B). Metabolic, cognitive, and gut microbiome outcomes were evaluated before and after a healthy dietary intervention (access to standard chow and water only) during early adulthood. PN: postnatal day, CAF: cafeteria diet, SUG: sugar diet, NOIC: novel object in context, Timepoint A: on western diet, Timepoint B: after dietary intervention, NMR: nuclear magnetic resonance imaging spectroscopy, IPGTT: intraperitoneal glucose tolerance test

### 2.3 Experiment 2 Design (Sugar-Sweetened Beverage)

At PN 26, rats were randomized to two groups matched for body weight and were given ad libitum access to standard chow and either: [1] 11% weight-by-volume (w/v) sugar-sweetened beverage solution (high fructose corn syrup type 55; Best Flavors, Orange, CA) diluted in water (SUG; n=16) or [2] a second bottle of water (CTL; n=16). Episodic memory was tested during young adulthood at PN 65 using the hippocampal-dependent NOIC task. At PN 67, rats were then tested for anxiety-like behavior using the Zero Maze. Fecal samples for microbiota analyses following sugar-sweetened beverage exposure (Timepoint A, n=16/group) were collected at PN 70. At PN 73, body composition using NMR was assessed. At PN 74, rodents underwent an IPGTT, and the dietary intervention where sugar solutions were removed was initiated. Post-dietary intervention body composition was measured at PN 132 and IPGTT was conducted at PN 134 (n=8/group; Figure 1C). A second cohort of juvenile (PN 25) female Sprague Dawley rats (n=18, 50-70g) were treated as described above for post dietary intervention behavioral testing. These rats were tested on NOIC after dietary intervention after 5 weeks of chow maintenance at PN 101 and Zero Maze at PN 107 (Figure 1D). Fecal samples for microbiome analysis were collected after dietary intervention at PN 105 (Timepoint B, n=9/group). Caloric intakes and body weights were recorded in this group as described above.

## General Procedures

### 2.4 Novel Object in Context

The Novel Object in Context (NOIC) task, which measures contextual episodic memory, was adapted from previous reports (Balderas et al., 2008; Martínez et al., 2014). Rats are habituated on consecutive days to both Context 1, a semi-transparent box (15in W × 24in L × 12in H) with yellow stripes, and Context 2, a grey opaque box (17in W × 17in L × 16in H) with the order counterbalanced between groups. Following the two habituation days, on the next day (Day 1 of NOIC), each animal is placed in Context 1 and allowed to explore two different objects (Object A and Object B) placed on diagonal, equidistant markings with ample space for the rat to circle the objects. The markings on which each object is placed is counterbalanced within each group. Objects used were either: a magic eight ball paired with a pyramid-shaped Lego object or a 12 oz. soda can (Coca-Cola®) paired with an upside-down stemless transparent wine glass. On the following day (NOIC day 2), rats are placed in Context 2 with Object A and a duplicate of Object A, one on each marking. On the subsequent day (test day; NOIC day 3), rats are placed again in Context 2 with Object A and Object B (which is not a novel object per se, but its presence in Context 2 is novel). Sessions are 5 minutes long and conducted under diffuse lighting conditions and are video recorded using an overhead camera (Digital USB 2.0 CMOS Camera with Vari-focal, 2.8-12mm lens; Stöelting) connected to a computer licensed with Any-Maze software (Stöelting, Wood Dale, IL), which objectively tracks the time spent investigating each object based on a programmed template. The discrimination index for the novel object is calculated as follows: (Time spent exploring Object B/ [Time spent exploring Object A + Time spent exploring Object B]). Intact rats typically preferentially explore Object B on NOIC day 3 due to its novelty in that particular context, an effect that is disrupted with hippocampal inactivation (Martínez et al., 2014), or by early life sugar consumption in male rats (Noble et al., 2019).

### 2.5 Zero Maze

The Zero Maze test was utilized to measure anxiety-like behavior (Shepherd et al., 1994). The maze consists of an elevated circular platform (63.5 cm height, 116.8cm external diameter) with two closed zones and two open zones, all of which are equal in length. The closed zones are enclosed with 17.5 cm high walls whereas the open zones have only 3 cm high curbs. Animals are placed in the maze for a single 5-min session, and the apparatus is cleaned with 10% ethanol in between animals. Any-Maze software scripts (Stoelting Co., Wood Dale, IL) are used to record videos and analyze the following parameters: time spent in the open zones and number of entries into the open zones.

### 2.4 Body Composition using NMR

Rats are food restricted for one hour prior to being weighed and scanned for body composition as previously described (Noble et al., 2017a) using the Bruker NMR Minispec LF 90II (Bruker Daltonics, Inc.), which is based on Time Domain NMR signals from all protons and has the benefit of being non-invasive and requiring no anesthesia. Percent body fat is calculated as [fat mass (g)/body weight (g)] x 100.

### 2.5 Intraperitoneal Glucose Tolerance Test

An intraperitoneal glucose tolerance test (IPGTT) was administered to estimate peripheral insulin sensitivity. Rats are food restricted for 22 hours prior to IPGTT. Baseline blood glucose readings are collected from blood sampled from the tip of the tail and measured using a glucometer (One touch Ultra2, LifeScan Inc., Milpitas, CA). Each rat receives an intraperitoneal injection of a 50% dextrose solution (0.5g/kg body weight) and blood glucose readings are obtained from the tail snip at the 0 (immediately before injection), 30, 60, 90, and 120 min timepoints following injection.

### 2.6 Fecal Sample Collection and 16s Ribosomal RNA (rRNA) Gene Sequencing

Rats were placed in a sterile cage with no bedding and were mildly restrained until defecation occurred. Two fecal samples were collected per animal. Samples were weighed under sterile conditions and then placed into a DNA/RNA free 2 mL cryogenic vial embedded in dry ice. Samples were then stored in a -80°C freezer until processing. The cages and all materials used to collect samples were cleaned with 70% ethanol in between rats. Bacterial genomic DNA was extracted from rat fecal samples using Laragen’s validated in-house fecal DNA extraction protocol. Quantification of 16S rRNA gene loads was performed by qPCR using the SYBR Green master mix (Roche), universal 16S rRNA gene primers55 and the QuantStudio 5 thermocycler (cycling parameters: 2 min at 50 °C, 10 min at 95 °C, 40 cycles of 10 s at 95 °C, and 45 s at 62 °C). Sequencing libraries were generated using methods previously described (Caporaso et al., 2011). The V4 regions of the 16S rRNA gene were PCR amplified using individually barcoded universal primers and 3 ng of the extracted genomic DNA. The PCR reaction was set up in a single reaction, and the PCR product was purified using Laragen’s validated in-house bead-based purification. Two hundred and fifty nanograms of purified PCR product from each sample was pooled and sequenced by Laragen, Inc. using the Illumina MiSeq platform and 2 × 150 bp reagent kit for paired-end sequencing.

### 2.7 Taxonomic Classification of 16S rRNA Gene Sequences

The sequencing reads were analyzed with QIIME2 and DADA2 following the developers’ instructions for quality control, de-noising and chimera removal (Callahan et al., 2016; Bolyen et al., 2019). The forward sequencing reads were truncated to 110 bp and denoised to 2067 amplicon sequence variants (ASVs) using DADA2. The chimeras were identified and removed using the DADA2 ‘consensus’ method. The ASVs were classified using the QIIME2 feature classifier classify-sklearn based on the SILVA database (release 132). The taxonomic abundance tables were normalized to correct for the different sequencing depth as previously described (Jones et al., 2018).

### 2.8 Statistics

Data, presented as means ± SEM, were analyzed and graphed using Prism software (GraphPad Inc., version 8.4.2), with significance set as p < 0.05. Body weights, caloric intake, and the IPGTT were analyzed using a Two-way mixed ANOVA with time as a within-subjects factor and diet as a between-subjects factor. Data were corrected for multiple comparisons using Sidak’s multiple comparison test. For the IPGTT, area under the curve (AUC) was measured using Prism software. The statistical analyses and visualization for microbiome outcomes were primarily performed using R statistical software (Version 3.6.3). The principal coordinates analysis (PCoA) of the rat gut microbiomes were calculated based on the Bray-Curtis dissimilarity at the genus level and visualized with functions in the R package ‘vegan’. The associations between gut microbiome and treatment were analyzed with the PERMANOVA test with 999 permutations using function ‘adonis’ in R package ‘vegan’. Shannon index was used for analyzing the alpha-diversity of gut microbiomes. The associations of individual taxa and treatment were analyzed with Wilcoxon test. The P-values were adjusted with the Benjamini-Hochberg method for multiple hypotheses testing. The correlations between each taxa and NOIC were analyzed with Spearman’s correlations and the P-values were adjusted with the Benjamini-Hochberg method.

## 3. Results

### 3.1 Increased adiposity associated with early life Western diet consumption is reversible with healthy dietary intervention

In Experiment 1, free access to the CAF diet throughout the juvenile and adolescence period did not result in significant differences in body weight (*F*_(1, 18)_ = 0.2917; *P* = 0.5958) or total caloric intake (*F*_(1, 18)_ = 2.065; *P* = 0.1679) (Fig. 2A-B). A significant time x diet interaction, however, was observed for total calories consumed (*F*_(33, 589)_ = 4.229; *P* < 0.0001). Surprisingly, this interaction was driven by reduced caloric consumption in the CAF group vs. controls following the healthy dietary intervention, as confirmed by a significant group main effect when analyzed separately over the post-intervention period (*F*_(1, 18)_ = 12.68; *P* = 0.0022) and the lack of significant group differences when analyzed separately over the CAF diet maintenance period before the intervention (*F*_(1, 18)_ = 0.1916; *P* = 0.6668). However, NMR-based body composition analyses revealed that CAF rats had elevated fat mass relative to controls at PN 74 after consuming a CAF during the juvenile and adolescence period (*P* = 0.0059). Importantly, the elevated adiposity in CAF vs. control group was normalized at PN 115 after 41 days of standard chow maintenance (Fig. 2E).

**Figure 2:**
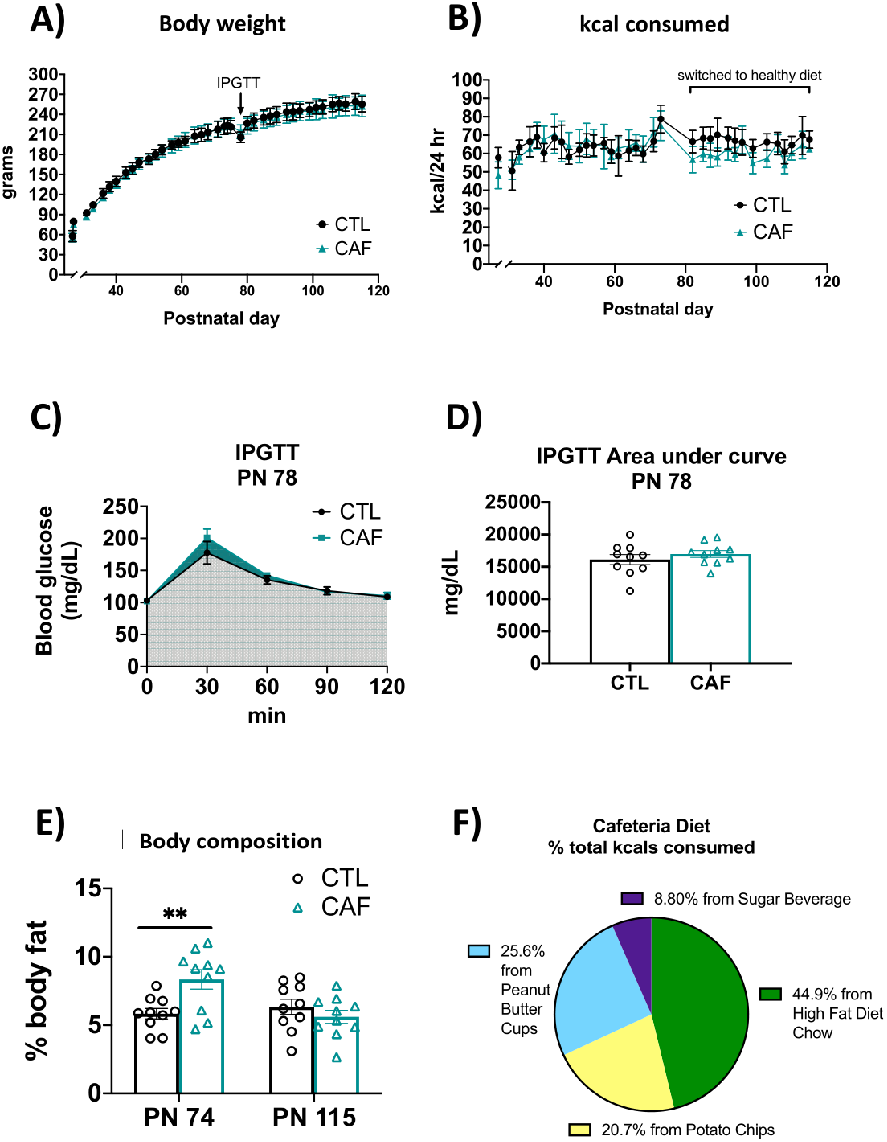
Energy balance and metabolic outcomes following adolescent cafeteria diet consumption. There were no overall significant group differences in body weight (A), total caloric intake (B), or glucose tolerance in the IPGTT test (C and D). Rats in the CAF group consumed significantly fewer calories following the healthy dietary intervention (B). CAF-exposed rats had significantly greater adiposity than CTL rats (E). Percent total calories from each food item in the CAF diet in Cohort 1 are depicted in (F). Data are means ± SEM; n = 10/group, **P < 0.01. CTL: control; CAF: cafeteria diet; kcal: kilocalories; IPGTT: intraperitoneal glucose tolerance test

Rats in the CAF group derived approximately 44.9% of their total calories from the high-fat-diet chow, 20.7% from potato chips, 25.6% from peanut butter cups, and 8.8% from the sugar beverage (Fig. 2F). All caloric intake and body weight data were comparable in the 2^nd^ cohort (Supplemental Figure 1).

In Experiment 2, body weights and total caloric intake were comparable in the SUG and control groups throughout the experiment from PN 26-137 despite *ad libitum* access to a sugar beverage from PN 26-74 in the SUG group (Fig. 4A and 4B). However, there was a significant interaction (time x diet) for calories consumed from standard chow (*F*_(39, 850)_ = 17.58; *P* < 0.0001) with a main effect of time (*F*_(7.808, 170.2) =_ 41.55; *P* < 0.0001) and diet (*F*_(1, 30)_ = 94.52; *P* < 0.0001). *Post hoc* analyses showed that from PN 28-73, SUG rats consumed fewer calories from chow (Fig. 4C), suggesting that the female SUG rats were reducing their chow intake to compensate for the kcals consumed from the sugar solution (Fig. 4D). Rats in the SUG group derived approximately 36.0% of their total calories from the sugar beverage (data not shown). Caloric intake and body weights within and between groups were comparable in the 2^nd^ cohort (Supplemental Figure 2), who also derived around 33.6% of their total calories from the sugar beverage (data not shown). Similarly, adiposity measured by body composition using NMR was higher in the SUG group before removal of the sugar at PN 73 (*P* = 0.0021), but not at PN 132 after dietary intervention (Fig. 2H).

### 3.2 Impaired peripheral glucose regulation induced by excessive early life sugar consumption is reversed by healthy dietary intervention

In Experiment 1, results from an IPGTT conducted before the CAF group was switched to chow maintenance revealed no significant group differences in area under the curve (AUC) (*P* = 0.3711) (Fig 2D), suggesting that glucose tolerance was not affected by CAF consumption. IPGTT results were comparable in the 2^nd^ cohort (Supplemental Figure 1). However, in Experiment 2, analyses of IPGTT results at PN 74 for the SUG and controls rats revealed a significant interaction (time x diet) for blood glucose levels (*F*_(4, 120)_ = 3.196, *P =* 0.0156) with a main effect of both time (*F*_(1.681, 50.42)_ = 231.8, *P* < 0.0001) and diet (*F*(_1, 30)_ = 7.642, *P* = 0.0097). *Post hoc* analyses revealed that the SUG group had higher glucose levels at the 120-minute timepoint after glucose administration (Fig. 2E). The IPGTT AUC analyses revealed that the SUG rats had a significantly higher glucose levels than CTL rats (*P* = 0.0106), further supporting that the SUG animals had impaired glucose tolerance at PN 74 (Fig. 2G). However, when re-tested at PN 137 after having removed the sugar for ∼1.5 months as a healthy dietary intervention, SUG rats did not exhibit significant glucose intolerance relative to controls (Fig. 2F).

### 3.3 Hippocampal-dependent memory impairments associated with early life junk food diet consumption are not reversible with healthy diet intervention

In experiment 1, results from the NOIC task revealed that CAF rats had deficits in hippocampal-dependent episodic contextual memory relative to controls, which was supported by a lower shift from baseline discrimination index for the novel object (P = 0.008; Fig. 3A). After switching the CAF animals to standard chow as a healthy dietary intervention, rats that had been exposed to the CAF diet during early life maintained a significantly reduced shift from baseline discrimination of the novel object (P = 0.02; Fig. 3B). This suggests that the memory impairment associated with ∼1.5 months of CAF diet exposure during adolescence was not reversible with ∼1.5 months of healthy dietary intervention. There were no differences in anxiety-like behavior in the Zero Maze test before switching to a low-fat diet at PN 67 or after the dietary intervention at PN 108 (Fig. 3C-D).

**Figure 3:**
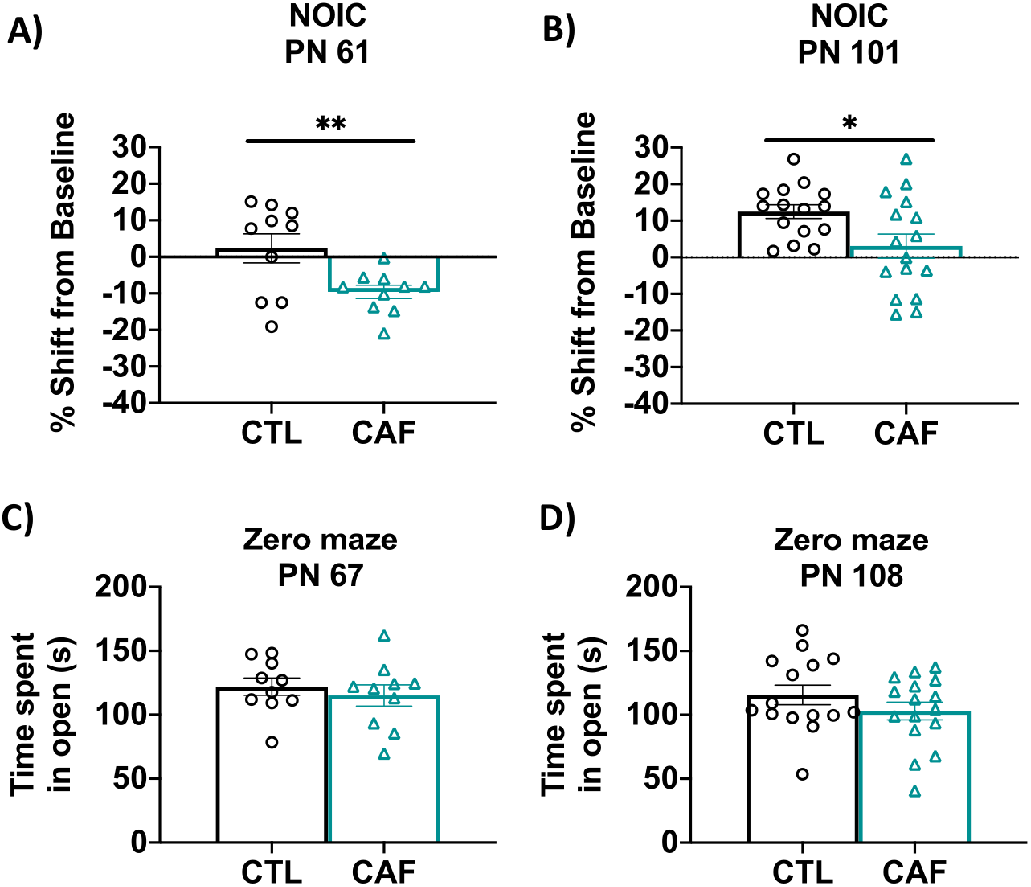
Hippocampal-dependent memory following adolescent cafeteria diet consumption. CAF-exposed rats were impaired in the NOIC memory task (calculated as shift from baseline discrimination index on test day) both before (A) and after (B) healthy dietary intervention. There were no significant group differences in anxiety-like behavior in the Zero Maze either before (C) or after (D) the dietary intervention. Data are means ± SEM; A/C: n = 10/group, B/D: n = 15/CTL group, n = 16/CAF group, *P < 0.05, **P < 0.01. CTL: control; CAF: cafeteria diet; NOIC = novel object in context

**Figure 4.**
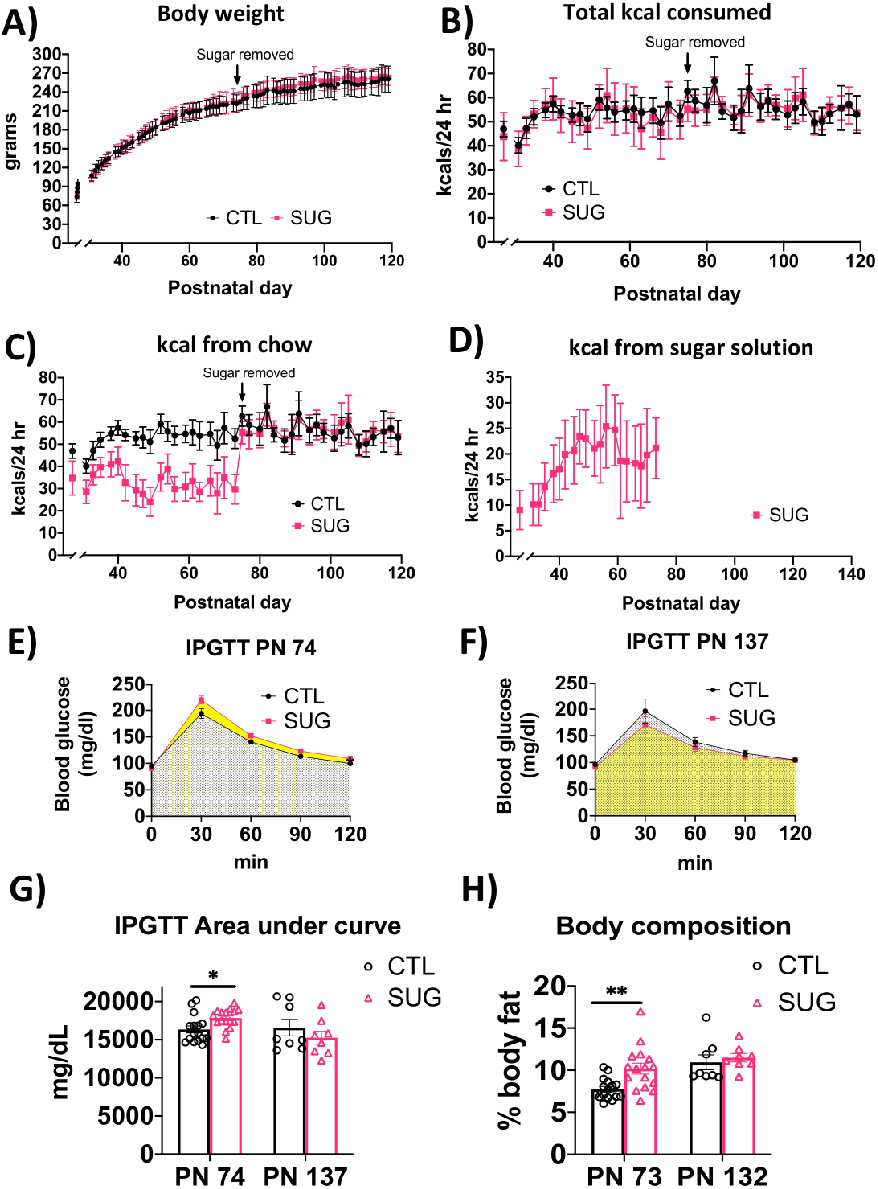
Energy balance and metabolic outcomes following adolescent sugar diet consumption. There were no overall significant group differences in body weight (A) or total caloric intake (B), although rats in the SUG group consumed less calories from chow (C) when consuming the sugar solution (D). The SUG group had significantly higher blood glucose levels (E and G) and adiposity (H) relative to CTL rats, which normalized after healthy dietary intervention (F, G and H). Data are means ± SEM; n=16/group before sugar removal (PN 74), n=8/group after sugar removal (PN 75+) *P < 0.05, **P < 0.01. CTL: control; SUG: sugar; kcal: kilocalories; IPGTT: intraperitoneal glucose tolerance test

In experiment 2, SUG rats were impaired in the hippocampal-dependent NOIC task relative to controls when tested at PN65, supported by a significantly lower shift from baseline discrimination index for the novel object relative to controls (*P* = 0.0309; Fig. 5A). After removing the SUG beverages as a dietary intervention, rats that had been previously exposed to the SUG diet during adolescence had a similar shift from baseline discrimination of the novel object as the CTL group (Fig. 5B). This suggests that the memory impairments associated with early life SUG consumption in female rats are reversible following ∼1.5 months of healthy dietary intervention. There were no differences in anxiety-like behavior in the Zero Maze test before removing the sugar beverage at PN 67 (Fig. 5C).

**Figure 5.**
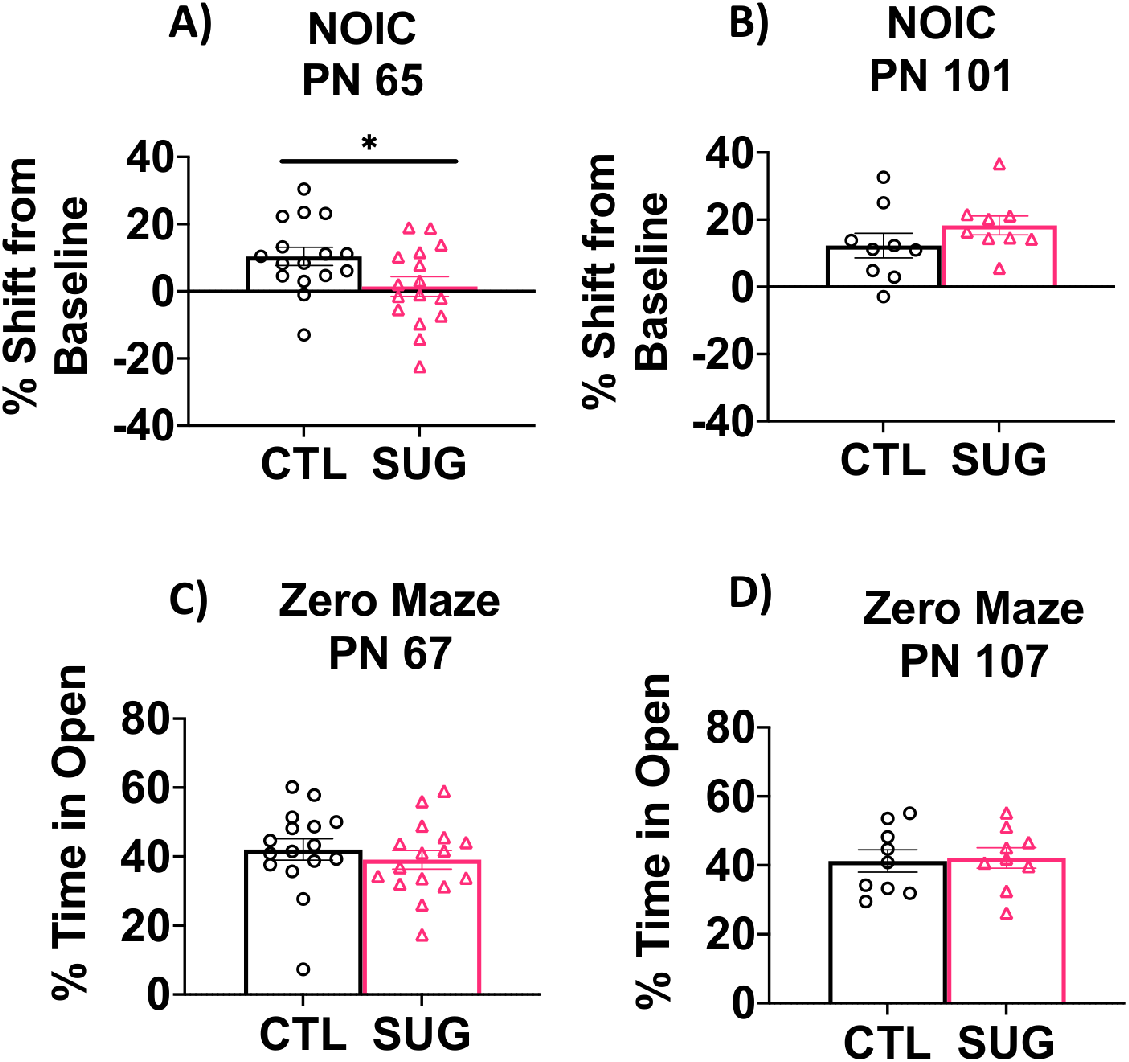
Hippocampal-dependent memory following adolescent sugar diet consumption. SUG-exposed rats were impaired in the NOIC memory task (calculated as shift from baseline discrimination index on test day) before (A), but not after (B) healthy dietary intervention. There were no significant group differences in anxiety-like behavior in the Zero Maze either before (C) or after (D) the dietary intervention. Data are means ± SEM; A/C: n=16/group, B/D: n=9/group *P < 0.05. CTL: control; SUG: sugar; NOIC = novel object in context

### 3.4 Early life Western diet consumption during adolescence produces gut microbiome changes that are not reversible following healthy dietary intervention

We characterized the rat gut microbiome to estimate the impact of early life Western diet consumption. The CAF and CTL groups’ gut microbiomes are separated in the PCoA plots for both timepoint A (PN70, 44 days after the start of the WD diet) and timepoint B (27 days after healthy dietary intervention), but the separation at timepoint B is stronger (Fig. 6A and B). The alpha diversity of CAF rats at timepoint B was significantly reduced compared to other groups (Fig. 6C).

**Figure 6.**
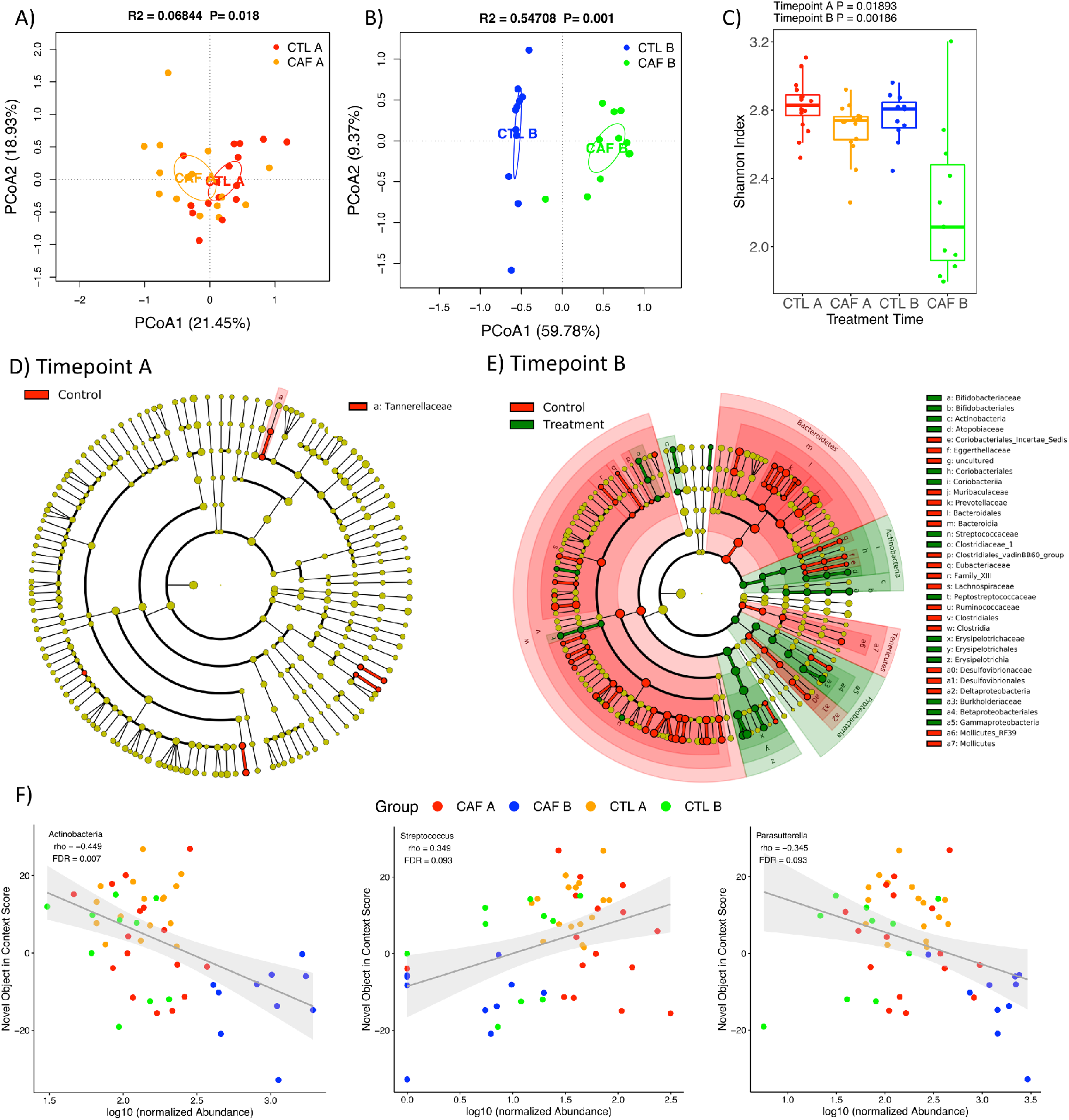
Gut microbiome following adolescent cafeteria diet consumption. (A and B) PCoA plots of CAF-exposed rats gut microbiomes before (A) and after (B) healthy dietary intervention. Ellipses indicate 95% confidence limits. R^2^ and P are from PERMANOVA tests. (C) Shannon index of CAF-exposed rats gut microbiomes before and after healthy dietary intervention. The differences were tested with Wilcoxon tests. (D and E) Significant differential taxa between treatment and control at phylum to species level (Wilcoxon test, FDR<0.05) are highlighted on the phylogenetic trees of all taxa identified in this study. An FDR cutoff of 0.05 was used here for visualization. (F) The correlations plots of Actinobacteria (phylum), *Streptococcus* and *Parasutterella* with NOIC performance across CAF and CTL samples. A complete list of significantly correlated taxa is shown in Table 1A. Spearman’s correlation was used for the analysis and P-values were corrected for multiple hypotheses testing with the Benjamini-Hochberg method. CTL A: control timepoint A; CAF A: cafeteria diet timepoint A; CTL B: control timepoint B; CAF B: cafeteria diet timepoint B; PCoA: Principal Coordinate Analysis; FDR: false discovery rate; rho: Spearman’s ρ

In order to generate a more detailed picture of changes to the microbiome, e next analyzed the differential abundance of individual taxa between CAF and CTL rats at timepoints A and B. Consistent with the view given by the PCoA plots, there are more taxa with significant group differences at timepoint B compared to timepoint A (151 taxa vs 9 taxa at class to genus level, Wilcoxon test, FDR<0.1; Fig. S4). The differential taxa at timepoint B are broadly distributed across various taxonomic groups, including Bacteroidetes and Clostridia that are more abundant in CTL rats and Gammaproteobacteria, Erysipelotrochia and Bifidobacteriales that are more abundant in CAF rats (Fig. 6D and E).

**Table 1.** The correlations between taxa and NOIC performance in CAF diet dataset (A) and SUG diet dataset (B). D_1 to D_6 indicate taxonomic levels from phylum to species. FDR: false discovery rate; rho: Spearman’s ρ

|  | All_rho | All_FDR | Treatment_rho | Treatment_FDR |
| --- | --- | --- | --- | --- |
| D_1__Actinobacteria | -0.449 | 0.007 | -0.549 | 0.034 |
| D_5__Ruminiclostridium.6 | 0.465 | 0.037 | 0.317 | 0.304 |
| D_5__Negativibacillus | 0.443 | 0.037 | 0.379 | 0.279 |
| D_5__Clostridium.sensu.stricto.1 | -0.419 | 0.049 | -0.332 | 0.279 |
| D_4__Family.XIII (Clostridiales) | 0.428 | 0.055 | 0.282 | 0.305 |
| D_2__Coriobacteriia | -0.386 | 0.065 | -0.433 | 0.091 |
| D_3__Coriobacteriales | -0.386 | 0.085 | -0.433 | 0.119 |
| D_5__Candidatus.Soleaferrea | 0.373 | 0.093 | 0.275 | 0.348 |
| D_5__Harryflintia | 0.364 | 0.093 | 0.357 | 0.279 |
| D_5__Lactococcus | -0.359 | 0.093 | -0.373 | 0.279 |
| D_5__Ruminococcaceae.UCG-005 | 0.356 | 0.093 | 0.198 | 0.531 |
| D_5__Streptococcus | 0.349 | 0.093 | 0.382 | 0.279 |
| D_5__Ruminiclostridium.9 | 0.344 | 0.093 | 0.342 | 0.279 |
| D_5__Parasutterella | -0.345 | 0.093 | -0.532 | 0.279 |
| D_5__Prevotellaceae.NK3B31.group | 0.333 | 0.099 | 0.372 | 0.279 |
| D_5__Holdemania | 0.332 | 0.099 | 0.353 | 0.279 |
| D_2__Erysipelotrichia | -0.323 | 0.130 | -0.475 | 0.091 |
| D_2__Actinobacteria | -0.285 | 0.135 | -0.447 | 0.091 |
| D_2__Gammaproteobacteria | -0.209 | 0.221 | -0.473 | 0.091 |
| D_1__Proteobacteria | -0.220 | 0.233 | -0.449 | 0.090 |

(B)
|  | All_rho | All_FDR | Treatment_rho | Treatment_FDR |
| --- | --- | --- | --- | --- |
| D_1__Actinobacteria | 0.230 | 0.180 | 0.428 | 0.091 |
| D_1__Bacteroidetes | -0.310 | 0.134 | -0.574 | 0.026 |
| D_1__Firmicutes | 0.266 | 0.134 | 0.451 | 0.091 |
| D_2__Bacteroidia | -0.310 | 0.218 | -0.574 | 0.041 |
| D_2__Actinobacteria | 0.296 | 0.218 | 0.502 | 0.075 |
| D_6__Alistipes.finegoldii | -0.316 | 0.188 | -0.418 | 0.093 |

Similar to Experiment 1, Experiment 2 PCoA plots also showed clear separation of the SUG and CTL rats’ gut microbiomes at both timepoints A and B, with a stronger separation at timepoint B (Fig. 7A and B). Unlike the CAF data, however, the alpha diversity of SUG and CTL rats was not significantly different for at either timepoint (Fig. 7C). Statistical tests of individual taxa abundance between SUG and CTL rats revealed 52 and 69 taxa that differed significantly at timepoints A and B, respectively (Wilcoxon test, FDR<0.1) (Fig. S5). The differential taxa at timepoint A include Actinobacteria and Gammaproteobacteria that are more abundant in SUG rats and Clostridia that is more abundant in CTL rats. The differential taxa at timepoint B include Actinobacteria and Erysipelotrochia that are more abundant in SUG rats and Bacteroidetes that is more abundant in CTL rats (Fig. 7D and E).

**Figure 7.**
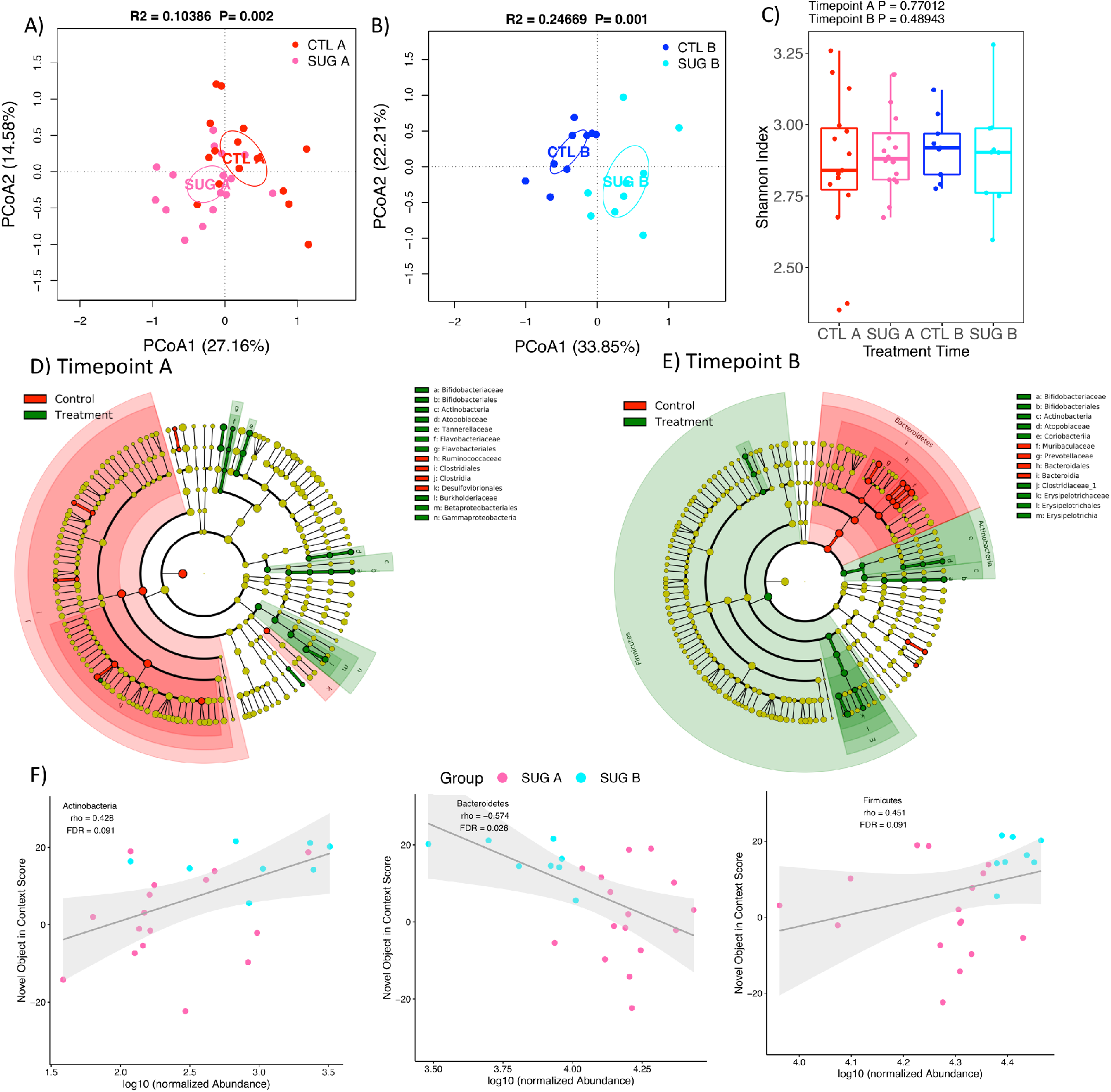
Gut microbiome following adolescent sugar diet consumption. (A and B) PCoA plots of SUG-exposed rats gut microbiomes before (A) and after (B) healthy dietary intervention. Ellipses indicate 95% confidence limits. R^2^ and P are from PERMANOVA tests. (C) Shannon index of SUG-exposed rats gut microbiomes before and after healthy dietary intervention. The differences were tested with Wilcoxon tests. (D and E) Significant differential taxa between treatment and control at phylum to species level (Wilcoxon test, FDR<0.05) are highlighted on the phylogenetic trees of all taxa identified in this study. An FDR cutoff of 0.05 was used here for visualization. (F) The correlations plots of Actinobacteria (phylum), Bacteroidetes and Firmicutes with NOIC performance across only SUG samples. A complete list of significantly correlated taxa is shown in Table 1B. Spearman’s correlation was used for the analysis and P-values were corrected for multiple hypotheses testing with the Benjamini-Hochberg method. CTL A: control timepoint A; CTL B: control timepoint B; SUG A: sugar timepoint A; SUG B: sugar timepoint B; PCoA: Principal Coordinate Analysis; FDR: false discovery rate; rho: Spearman’s ρ

PCoA plots also showed separation in the CTL rats when comparing timepoints A and B (Fig. S3A and C). It is not uncommon for microbial communities to change with time independent of experimental manipulations in animal facilities (Fan et al., 2018), but in our cohort changes associated with time in the control group, while marginally significant, are substantially smaller in effect size than changes associated both with diet and time (Fig. S3B and D).These outcomes indicate that the WD treatments induced a substantial change in the microbial community that was not reversible with healthy dietary intervention.

### 3.5 Microbiome changes are associated with changes in NOIC

We next analyzed the correlations between individual taxa and NOIC to explore the potential microbiome signature associated with memory performance. In the CAF diet study, we found many taxa significantly correlated with NOIC (Spearman’s correlation, FDR<0.1) (Table 1A, Fig 6F). Actinobacteria (phylum), *Clostridium sensu stricto 1*, Coriobacteriia, Coriobacteriales, *Lactococcus* and *Parasutterella* were negatively correlated with NOIC, while *Ruminiclostridium 6, Negativibacillus*, Clostridiales Family XIII, Candidatus *Soleaferrea, Harryflintia*, Ruminococcaceae UCG-005, *Streptococcus, Ruminiclostridium 9, Prevotellaceae NK3B31 group* and *Holdemania* were positively correlated. Actinobacteria (class), Erysipelotrichia, Gammaproteobacteria and Proteobacteria are correlated with NOIC only in the treatment group. In the SUG experiment (Experiment 2), while there were no taxa significantly correlated with NOIC across all the samples, taxa including Actinobacteria (phylum), Bacteroidetes, Firmicutes, Bacteroidia, Actinobacteria (class) and *Alistipes finegoldii* are significantly correlated with NOIC performance in the SUG group (Table 1B, Fig. 7F). Scatter plots of all taxa significantly correlated with NOIC are shown in Fig. S6. However, when adjusted for both treatment and timepoints, the associations are no longer significant, indicating that the correlations identified could be confound by the groups. Future studies are needed to unravel the complex interactions between WD, gut microbiome and NOIC performance.

## 4. Discussion

Our results reveal that adult female rats who were maintained on a CAF diet throughout adolescence demonstrate impaired HPC-dependent episodic memory, altered abundances of gut microbiome bacterial taxa, and increased adiposity relative to controls maintained on a healthy standard rodent diet. The CAF diet had no effect, however, on total caloric intake, body weight, or glucose tolerance relative to controls. Similar to previous results in male rats (Hsu et al., 2015b; Noble et al., 2019), our results further reveal that adolescent access to healthy chow, water, and an 11% carbohydrate w/v sugar solution (SUG) in female rats impaired HPC-dependent episodic memory, altered the gut microbiome, increased adiposity, and impaired glucose tolerance without affecting body weight or total caloric intake relative to controls. Additional results revealed that some, but not all of these early life WD-associated outcomes were reversible following a 5-week healthy diet intervention in which all animals were maintained on a standard control rodent diet beginning at early adulthood. For the CAF rats, the dietary intervention did not reverse the episodic memory deficits or the separation in gut microbial taxa, whereas the increased adiposity relative to controls normalized. For the SUG rats, all metabolic and cognitive outcomes were reversed by dietary intervention. However, similar to the CAF rats, the gut microbiome from SUG rats remained distinct from controls despite the dietary intervention. Together, these results show that some, but not all, negative outcomes associated with early life Western diet (WD) consumption in female rats are reversible by switching to a healthy diet.

Consistent with the present results, previous studies conducted in male rodents have also shown that dietary intervention ameliorates metabolic deficits associated with WD consumption (high fat or high fat, high sugar diets) (Guo et al., 2009; South et al., 2012; Kowalski et al., 2016; Hatzidis et al., 2017; Crisóstomo et al., 2019; Noble et al., 2019). However, some of the present results relevant to WD-associated memory outcomes differ from studies conducted in male rodents. For instance, unlike the female CAF rats in the present study, switching to a low fat diet following high fat diet adolescent exposure in male rats resulted in marginal improvements in HPC-dependent memory (Boitard et al., 2016) and cognitive flexibility (McNeilly et al., 2016). In another study, male rats that received an adolescent CAF diet were not impaired in a spatial memory task despite showing symptoms of metabolic syndrome, which were reversible by switching to standard chow (Gomez-Smith et al., 2016). Moreover, studies using a diet manipulation similar to our SUG model revealed that a high sugar diet during adolescence led to impairments in both HPC-dependent episodic (Noble et al., 2019) and spatial memory (Fierros-Campuzano et al., 2020) in male rats that were not reversible by dietary intervention (i.e., removing the sugar access). Compared to these findings in males, the present study overall suggests that female rats may benefit more from dietary intervention for neurocognitive impairments following early life excessive sugar consumption, despite consuming more of their total calories from the sugar beverage than males (here, the females consumed around ∼33-36% of their daily energy intake from the sugar beverage whereas Noble et al. found that males on the same diet consumed on average ∼24% of their daily energy intake from the sugar beverage) (Noble et al., 2019). There is also evidence to suggest that even metabolic and place-recognition memory impairments due to excessive sugar consumption by female rats in adulthood can be reversed with dietary intervention (switching to saccharin or water) (Kendig et al., 2018). However, female rats may be more susceptible to sustained detrimental effects on memory when the diet is high in both sugar and fat (e.g., the CAF diet). Future work is needed to disentangle the role of sex and sex hormones on the enduring neurocognitive outcomes associated with early life WD consumption.

Given that increased anxiety-like behavior may develop following early life WD exposure in rodents (Tsan et al., 2021), we tested the rats in the Zero Maze test but found no effects of CAF exposure on anxiety-like behavior either before or after the dietary intervention. Similar to our CAF rats, an anxiety-like phenotype was not seen in the SUG cohorts, which is consistent with our previous findings in male rats using the Zero Maze test (Hsu et al., 2015b; Noble et al., 2019). However, other studies have reported an anxiety-like phenotype in the open field test in male rodents after switching to standard chow for 1 week following an adolescent CAF diet (Lalanza et al., 2014) and after adolescent SUG exposure even after removing the sugar access for 25 days in young adulthood (Kruse et al., 2019). Thus, it is possible that the detection of anxiety-like behavior after WD exposure may be more or less sensitive depending on the type of behavioral test used.

Our data show that gut microbial richness, as measured with the Shannon Index, is significantly reduced following early life CAF, but not SUG, diet, an outcome that was actually exacerbated following the healthy dietary intervention. This outcome is consistent with other studies in which microbial richness was reduced after the removal of high-fat, high-sugar diets (Fülling et al., 2020; McNamara et al., 2021) or reduced adiposity following lifestyle modifications (eating breakfast, avoiding sugar-sweetened beverages, decreasing processed foods rich in animal fat or lengthening meal duration, and implementing more exercise) (Cho, 2021). Composition of the gut microbiota was significantly distinct from controls immediately after either the adolescent CAF or SUG exposure period analyzed with PERMANOVA tests. Surprisingly, the microbial separation between CAF and controls and between SUG and controls was greater after the 5-week healthy dietary intervention, suggesting that diet-induced shifts in the gut microbiome that occur during early life periods of development may be long-lasting independent of dietary patterns during adulthood. The comparison of two timepoints within each group indicates that changes to the microbiome in the animal facility or animal life stage associated with time had a significant influence on the gut microbiome as well. However, this shift in the control group is not as large as that in the treatment group possibly because of the healthy dietary intervention (Fig. S3). However, the healthy diet during adulthood did not shift the gut microbiome in the treatment towards that of the control group that was fed a healthy diet since adolescence.

In some cases, the same bacterial taxa were altered by both CAF and SUG treatment, but in opposite directions, and differential by timepoint. For example, at timepoint A (before dietary intervention), the CAF and SUG group shared changes in the class Gammaproteobacteria, the order Betaproteobacteriales, the family *Tannerellaceae*, and the genera *Parabacteroides* and *ASF356*, yet all of these followed the same trend and differed by dietary treatment: the abundances of these taxa were lower in CAF rats vs. controls and higher in SUG rats vs. controls. Looking at these same taxonomic levels at timepoint B, abundances in the class Gammaproteobacteria, which is more abundant in obese mice (Pindjakova et al., 2017) and increased after consumption of high-fructose syrup in honeybees (Wang et al., 2020), and the order Betaproteobacteriales were still significantly different in the CAF, but not SUG, group relative to their respective controls, but the direction changed: the values of these taxa were lower than controls in CAF rats prior to intervention, yet higher than controls after dietary intervention in the CAF-treated animals. At timepoint B, the class Mollicutes and the order Mollicutes RF39 were the only taxa that were significantly different in both CAF and SUG treated groups, lower in the CAF group, but higher in the SUG group compared to respective controls. Overall, it is clear that CAF and SUG diets both significantly altered the gut microbiomes, but the changes in microbiota were often divergent. These dietary treatments had significant and distinct influences on the microbiomes that persisted after the healthy dietary intervention.

There is evidence to suggest that gut dysbiosis occurs before metabolic and spatial memory impairment in rodents (Saiyasit et al., 2020) and that certain bacteria can improve memory performance (Romo-Araiza et al., 2018; Ishikawa et al., 2019) or have detrimental effects on HPC neurons (Li et al., 2019) and cognitive ability (Magnusson et al., 2015). Further, we recently showed that elevated levels of *Parabacteroides* in males rats given the SUG dietary treatment during adolescence were functionally connected to HPC-dependent memory impairments (Noble et al., 2021). This conclusion was based on findings that levels of *Parabacteroides* were negatively associated with memory performance, and targeted enrichment of *Parabacteroides* in rats that had never consumed sugar replicated the memory deficits. Similar to these findings, here we show in female rats that *Parabacteroides* levels were increased in the SUG group relative to controls, and interestingly, that this elevation reversed following the dietary intervention. Given that the HPC-dependent memory impairments were also reversed by dietary intervention in this group, this suggests that levels *Parabacteroides* may be functionally connected to SUG-associated memory impairments in females as well. However, correlation analyses revealed that *Parabacteroides* abundance was not significantly correlated with memory performance in females after adjusting for multiple hypotheses testing, suggesting the associations between *Parabacteroides* and SUG-associated memory impairments may vary by the sex of rats. It is also possible that this association in female rats may require a larger sample size than male rats for adequate statistical power.

Our data identify multiple taxa that are significantly correlated with NOIC performance in CAF-treated animals and CTLs, including *Negativibacillus*, Clostridiales family *Family XIII, Prevotellaceae NK3B31 group, Ruminiclostridium 6, Harryflintia*, and *Candidatus Soleaferrea*. This may be related to the poor performance in the NOIC task in the CAF group at timepoint B, where most of these taxa were found to be significantly higher or lower in abundance. While there were no taxa significantly correlated with NOIC in the SUG-treated rats and controls, Actinobacteria (phylum), Bacteroidetes, Firmicutes, Bacteroidia, Actinobacteria (class) and *Alistipes finegoldii* are significantly correlated with NOIC performance in the treatment group where the impairment was reversed by dietary intervention (timepoint B). Collectively, these data highlight various bacterial taxa of significant associations with the enduring poor memory performance seen in the NOIC task following early life CAF exposure. However, the associations between these taxa and NOIC were confounded by the treatment and timepoint groups, and future studies will be required to determine the causal relationships between these bacteria and NOIC performance.

In contrast to our results, some studies have reported that dietary interventions involving a switch to healthier diets following WD can alleviate gut dysbiosis in rodents (Zhang et al., 2012; Shang et al., 2017; Safari et al., 2019) and in humans (Haro et al., 2017; Qian et al., 2020). Our results, however, show an even greater divergence in the microbiome relative to controls after switching from either a SUG or a CAF diet to a healthy control diet for ∼5 weeks, especially in the CAF animals. These discrepancies could be due to the timing of when WD was introduced (we exposed animals to the WD during a critical developmental period, whereas the studies cited above evaluated WD consumption in adulthood). Thus, the present results reinforce that diet composition during early life is critical to the composition of the gut microbiome in adulthood. Although further research needs to be conducted, this divergence despite healthy dietary intervention may be related to altered competition within the gut microbial community due to the long-term effects of an early life WD. Understanding the influence of early life diets on the competitive landscape of the gut microbiome will be key in helping to reverse WD-induced gut dysbiosis and its effect on cognitive ability. One possible therapy to explore further is the use of probiotics, especially since there is evidence to suggest that probiotics can help improve memory function after WD exposure in male rodents (Beilharz et al., 2018; Chunchai et al., 2018; Yang et al., 2019) and help improve WD-induced gut dysbiosis in female mice (Kong et al., 2019).

In conclusion, we found that consumption of either a high sugar diet or a cafeteria-style junk food diet during adolescence can lead to metabolic dysfunction, HPC-dependent episodic memory deficits, and gut microbiome dysbiosis in female rats. The metabolic outcomes can be reversed by a 5-week healthy dietary intervention (chow and water only), whereas HPC-dependent memory deficits are reversible following adolescent excessive sugar consumption, but not following the junk food diet maintenance. The changes in the gut microbiome relative to controls, however, were not reversible with dietary intervention, suggesting that dietary effects on gut bacterial populations during early life periods may have long-lasting implications. In the CAF group, we identified several taxa whose abundance was altered by diet and correlated with performance in the HPC-dependent memory task. Cross study comparisons of present results with the literature identify the need to consider sex as a key variable in studying connections between WD, the microbiome, and neurocognitive outcomes.

**Supplemental Figure 1.**
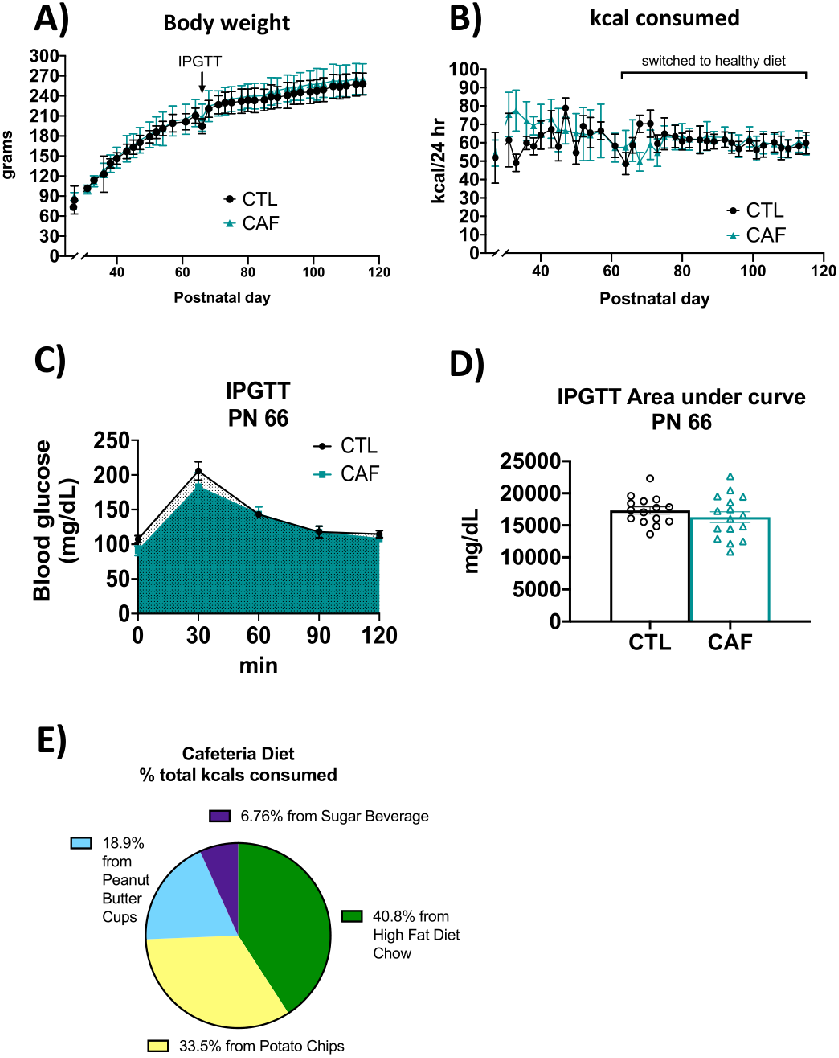
Energy balance and metabolic outcomes following adolescent cafeteria diet consumption in CAF cohort 2. There were no overall significant group differences in body weight (A), total caloric intake (B), or glucose tolerance in the IPGTT test (C and D), although rats in the CAF group consumed significantly fewer calories following the healthy dietary intervention (B, right). Percent total calories from each food item in the CAF diet in cohort 2 are depicted in (E). Data are means ± SEM; CTL: control n=15; CAF: cafeteria diet n=16; kcal: kilocalories; IPGTT: intraperitoneal glucose tolerance test

**Supplemental Figure 2.**
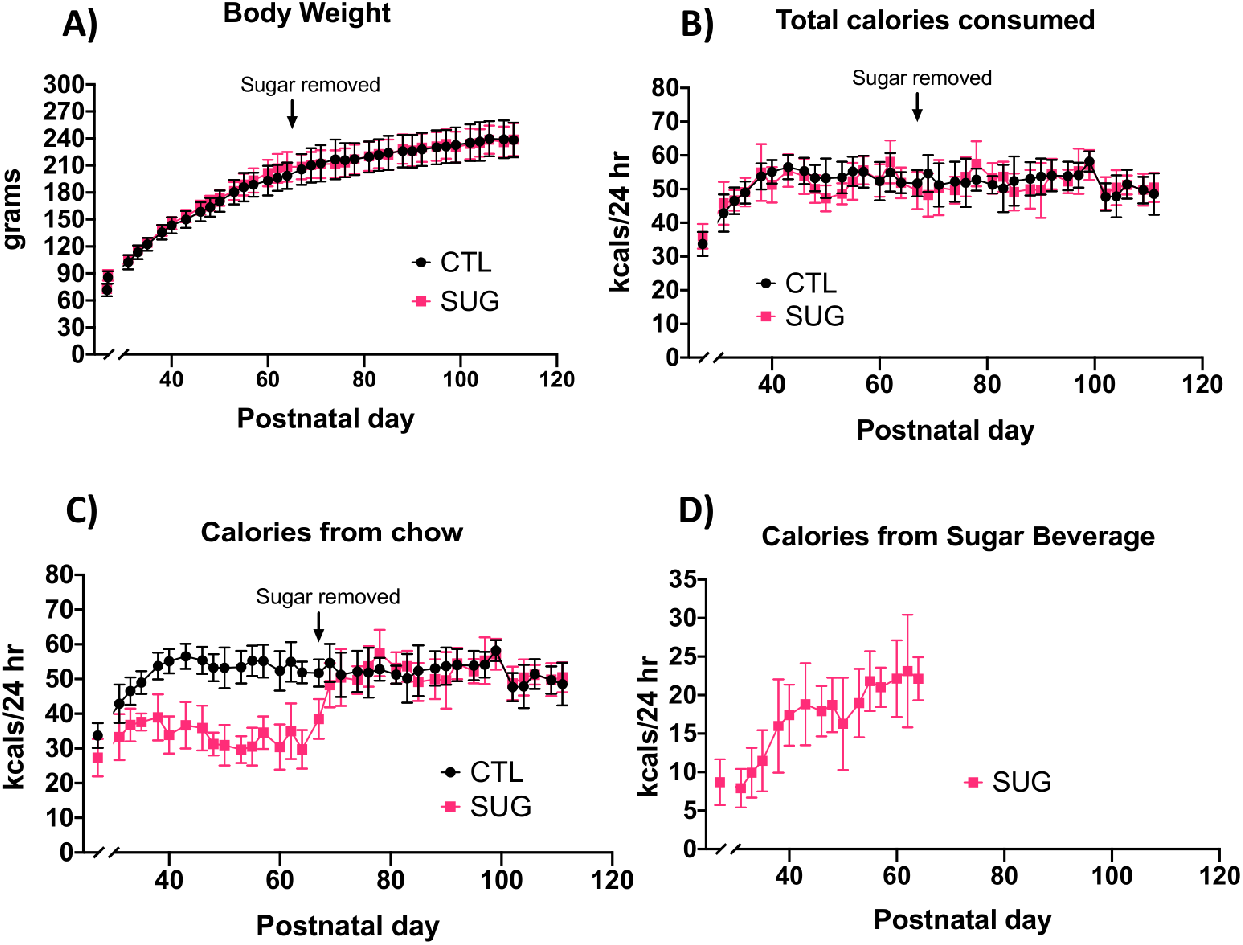
Energy balance and metabolic outcomes following adolescent sugar diet consumption in SUG cohort 2. There were no overall significant group differences in body weight (A) or total caloric intake (B), although rats in the SUG group consumed less calories from chow (C) when consuming the sugar solution (D). Data are means ± SEM; n=9/group CTL: control; SUG: sugar; kcal: kilocalories

**Supplemental Figure 3.**
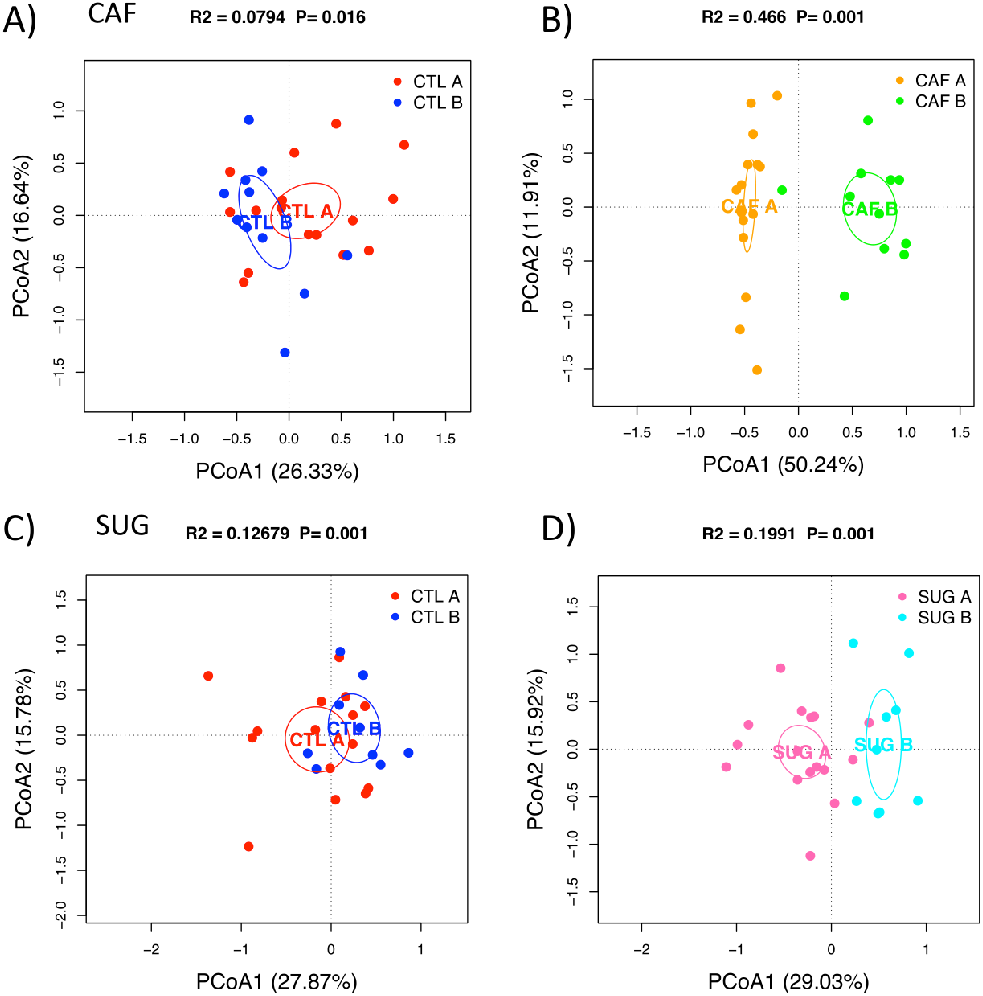
Additional PCoA of the gut microbiome by group. There was significant separation between the CTL groups at timepoint A (before dietary intervention) and B (after dietary intervention) for the CAF experiment (A) and the SUG experiment (C). There was also significant separation between the CAF groups at timepoint A and B (B) and the SUG groups at timepoint A and B (D). Ellipses indicate 95% confidence limits. R^2^ and P are from PERMANOVA tests. CTL: control; CAF: cafeteria diet; SUG: sugar; PCoA: Principal Coordinate Analysis

**Supplemental Figure 4 Abundance of the gut microbiota in the CAF vs CTL groups before and after healthy dietary intervention**. CTL A: control timepoint A; CAF A: cafeteria diet timepoint A; CTL B: control timepoint B; CAF B: cafeteria diet timepoint B; Café: cafeteria diet; FDR: false discovery rate

**Supplemental Figure 5 Abundance of the gut microbiota in the SUG vs CTL groups before and after healthy dietary intervention**. CTL A: control timepoint A; SUG A: sugar timepoint A; CTL B: control timepoint B; SUG B: sugar timepoint B; FDR: false discovery rate

**Supplemental Figure 6 Significant correlations with NOIC performance in the CAF experiment and SUG experiment**. Spearman’s correlation was used for the analysis and P-values were corrected for multiple hypotheses testing with the Benjamini-Hochberg method. CTL A: control timepoint A; CAF A: cafeteria diet timepoint A; CTL B: control timepoint B; CAF B: cafeteria diet timepoint B; SUG A: sugar timepoint A; SUG B: sugar timepoint B; FDR: false discovery rate; rho: Spearman’s ρ

## Acknowledgments

This work was supported by grant numbers DK116942 (Awarded to SK and AF) from the National Institute of Diabetes and Digestive and Kidney Diseases and also by the National Science Foundation Graduate Research Fellowship DGE-1842487 (Awarded to LT).

## References

Ambrosini, G. L., Huang, R.-C., Mori, T. A., Hands, B. P., O’Sullivan, T. A., de Klerk, N. H., et al. (2010). Dietary patterns and markers for the metabolic syndrome in Australian adolescents. Nutrition, Metabolism and Cardiovascular Diseases 20, 274–283. doi:10.1016/j.numecd.2009.03.024.

Balderas, I., Rodriguez-Ortiz, C. J., Salgado-Tonda, P., Chavez-Hurtado, J., McGaugh, J. L., and Bermudez-Rattoni, F. (2008). The consolidation of object and context recognition memory involve different regions of the temporal lobe. Learn. Mem. 15, 618–624. doi:10.1101/lm.1028008.

Baym, C. L., Khan, N. A., Monti, J. M., Raine, L. B., Drollette, E. S., Moore, R. D., et al. (2014). Dietary lipids are differentially associated with hippocampal-dependent relational memory in prepubescent children. Am J Clin Nutr 99, 1026–1032. doi:10.3945/ajcn.113.079624.

Beilharz, J. E., Kaakoush, N. O., Maniam, J., and Morris, M. J. (2018). Cafeteria diet and probiotic therapy: cross talk among memory, neuroplasticity, serotonin receptors and gut microbiota in the rat. Molecular Psychiatry 23, 351–361. doi:10.1038/mp.2017.38.

Boitard, C., Parkes, S. L., Cavaroc, A., Tantot, F., Castanon, N., Layé, S., et al. (2016). Switching Adolescent High-Fat Diet to Adult Control Diet Restores Neurocognitive Alterations. Front Behav Neurosci 10. doi:10.3389/fnbeh.2016.00225.

Bolyen, E., Rideout, J. R., Dillon, M. R., Bokulich, N. A., Abnet, C. C., Al-Ghalith, G. A., et al. (2019). Reproducible, interactive, scalable and extensible microbiome data science using QIIME 2. Nat Biotechnol 37, 852–857. doi:10.1038/s41587-019-0209-9.

Callahan, B. J., McMurdie, P. J., Rosen, M. J., Han, A. W., Johnson, A. J. A., and Holmes, S. P. (2016). DADA2: High-resolution sample inference from Illumina amplicon data. Nature Methods 13, 581–583. doi:10.1038/nmeth.3869.

Caporaso, J. G., Lauber, C. L., Walters, W. A., Berg-Lyons, D., Lozupone, C. A., Turnbaugh, P. J., et al. (2011). Global patterns of 16S rRNA diversity at a depth of millions of sequences per sample. Proc Natl Acad Sci U S A 108, 4516–4522. doi:10.1073/pnas.1000080107.

Cerdó, T., Diéguez, E., and Campoy, C. (2019). Early nutrition and gut microbiome: interrelationship between bacterial metabolism, immune system, brain structure, and neurodevelopment. American Journal of Physiology-Endocrinology and Metabolism 317, E617–E630. doi:10.1152/ajpendo.00188.2019.

Cho, K. Y. (2021). Lifestyle modifications result in alterations in the gut microbiota in obese children. BMC Microbiology 21, 10. doi:10.1186/s12866-020-02002-3.

Chunchai, T., Thunapong, W., Yasom, S., Wanchai, K., Eaimworawuthikul, S., Metzler, G., et al. (2018). Decreased microglial activation through gut-brain axis by prebiotics, probiotics, or synbiotics effectively restored cognitive function in obese-insulin resistant rats. J Neuroinflammation 15, 11. doi:10.1186/s12974-018-1055-2.

Clark, K. A., Alves, J. M., Jones, S., Yunker, A. G., Luo, S., Cabeen, R. P., et al. (2020). Dietary Fructose Intake and Hippocampal Structure and Connectivity during Childhood. Nutrients 12, 909. doi:10.3390/nu12040909.

Costa, C. S., Del-Ponte, B., Assunção, M. C. F., and Santos, I. S. (2018). Consumption of ultra-processed foods and body fat during childhood and adolescence: a systematic review. Public Health Nutrition 21, 148–159. doi:10.1017/S1368980017001331.

Crisóstomo, L., Rato, L., Jarak, I., Silva, B. M., Raposo, J. F., Batterham, R. L., et al. (2019). A switch from high-fat to normal diet does not restore sperm quality but prevents metabolic syndrome. Reproduction 158, 377–387. doi:10.1530/REP-19-0259.

Darch, H. T., Collins, M. K., O’Riordan, K. J., and Cryan, J. F. (2021). Microbial memories: Sex-dependent impact of the gut microbiome on hippocampal plasticity. Eur J Neurosci. doi:10.1111/ejn.15119.

Fan, T.-J., Tchaptchet, S. Y., Arsene, D., Mishima, Y., Liu, B., Sartor, R. B., et al. (2018). Environmental Factors Modify the Severity of Acute DSS Colitis in Caspase-11-Deficient Mice. Inflamm Bowel Dis 24, 2394–2403. doi:10.1093/ibd/izy244.

Ferreira, A., Castro, J. P., Andrade, J. P., Dulce Madeira, M., and Cardoso, A. (2018). Cafeteria-diet effects on cognitive functions, anxiety, fear response and neurogenesis in the juvenile rat. Neurobiol Learn Mem 155, 197–207. doi:10.1016/j.nlm.2018.07.014.

Fierros-Campuzano, J., Ballesteros-Zebadúa, P., Manjarrez-Marmolejo, J., Aguilera, P., Méndez-Diaz, M., Prospero-García, O., et al. (2020). Irreversible hippocampal changes induced by high fructose diet in rats. Nutr Neurosci, 1–13. doi:10.1080/1028415X.2020.1853418.

Francis, H., and Stevenson, R. (2013). The longer-term impacts of Western diet on human cognition and the brain. Appetite 63, 119–128. doi:10.1016/j.appet.2012.12.018.

Fülling, C., Lach, G., Bastiaanssen, T. F. S., Fouhy, F., O’Donovan, A. N., Ventura-Silva, A.-P., et al. (2020). Adolescent dietary manipulations differentially affect gut microbiota composition and amygdala neuroimmune gene expression in male mice in adulthood. Brain, Behavior, and Immunity. doi:10.1016/j.bbi.2020.02.013.

Gallagher, C., Moschonis, G., Lambert, K. A., Karaglani, E., Mavrogianni, C., Gavrili, S., et al. (2020). Sugar-sweetened beverage consumption is associated with visceral fat in children. Br J Nutr, 1–9. doi:10.1017/S0007114520003256.

Gomez-Smith, M., Karthikeyan, S., Jeffers, M. S., Janik, R., Thomason, L. A., Stefanovic, B., et al. (2016). A physiological characterization of the Cafeteria diet model of metabolic syndrome in the rat. Physiology & Behavior 167, 382–391. doi:10.1016/j.physbeh.2016.09.029.

Guo, J., Jou, W., Gavrilova, O., and Hall, K. D. (2009). Persistent Diet-Induced Obesity in Male C57BL/6 Mice Resulting from Temporary Obesigenic Diets. PLoS One 4. doi:10.1371/journal.pone.0005370.

Haro, C., García-Carpintero, S., Rangel-Zúñiga, O. A., Alcalá-Díaz, J. F., Landa, B. B., Clemente, J. C., et al. (2017). Consumption of Two Healthy Dietary Patterns Restored Microbiota Dysbiosis in Obese Patients with Metabolic Dysfunction. Molecular Nutrition & Food Research 61, 1700300. doi:https://doi.org/10.1002/mnfr.201700300.

Hatzidis, A., Hicks, J. A., Gelineau, R. R., Arruda, N. L., De Pina, I. M., O’Connell, K. E., et al. (2017). Removal of a high-fat diet, but not voluntary exercise, reverses obesity and diabetic-like symptoms in male C57BL/6J mice. Hormones 16, 62–74. doi:10.14310/horm.2002.1720.

Hsu, T. M., Konanur, V. R., Taing, L., Usui, R., Kayser, B. D., Goran, M. I., et al. (2015a). Effects of sucrose and high fructose corn syrup consumption on spatial memory function and hippocampal neuroinflammation in adolescent rats. Hippocampus 25, 227–239. doi:10.1002/hipo.22368.

Hsu, T. M., Konanur, V. R., Taing, L., Usui, R., Kayser, B. D., Goran, M. I., et al. (2015b). Effects of sucrose and high fructose corn syrup consumption on spatial memory function and hippocampal neuroinflammation in adolescent rats. Hippocampus 25, 227–239. doi:10.1002/hipo.22368.

Ishikawa, R., Fukushima, H., Nakakita, Y., Kado, H., and Kida, S. (2019). Dietary heat-killed Lactobacillus brevis SBC8803 (SBL88TM) improves hippocampus-dependent memory performance and adult hippocampal neurogenesis. Neuropsychopharmacology Reports 39, 140–145. doi:https://doi.org/10.1002/npr2.12054.

Jones, R. B., Zhu, X., Moan, E., Murff, H. J., Ness, R. M., Seidner, D. L., et al. (2018). Inter-niche and inter-individual variation in gut microbial community assessment using stool, rectal swab, and mucosal samples. Sci Rep 8, 4139. doi:10.1038/s41598-018-22408-4.

Kashtanova, D. A., Popenko, A. S., Tkacheva, O. N., Tyakht, A. B., Alexeev, D. G., and Boytsov, S. A. (2016). Association between the gut microbiota and diet: Fetal life, early childhood, and further life. Nutrition 32, 620–627. doi:10.1016/j.nut.2015.12.037.

Kendig, M. D., Boakes, R. A., Rooney, K. B., and Corbit, L. H. (2013). Chronic restricted access to 10% sucrose solution in adolescent and young adult rats impairs spatial memory and alters sensitivity to outcome devaluation. Physiology & Behavior 120, 164–172. doi:10.1016/j.physbeh.2013.08.012.

Kendig, M. D., Fu, M. X., Rehn, S., Martire, S. I., Boakes, R. A., and Rooney, K. B. (2018). Metabolic and cognitive improvement from switching to saccharin or water following chronic consumption by female rats of 10% sucrose solution. Physiology & Behavior 188, 162–172. doi:10.1016/j.physbeh.2018.02.008.

Kong, C., Gao, R., Yan, X., Huang, L., and Qin, H. (2019). Probiotics improve gut microbiota dysbiosis in obese mice fed a high-fat or high-sucrose diet. Nutrition 60, 175–184. doi:10.1016/j.nut.2018.10.002.

Kowalski, G. M., Hamley, S., Selathurai, A., Kloehn, J., De Souza, D. P., O’Callaghan, S., et al. (2016). Reversing diet-induced metabolic dysregulation by diet switching leads to altered hepatic de novo lipogenesis and glycerolipid synthesis. Sci Rep 6. doi:10.1038/srep27541.

Kruse, M. S., Vadillo, M. J., Miguelez Fernández, A. M. M., Rey, M., Zanutto, B. S., and Coirini, H. (2019). Sucrose exposure in juvenile rats produces long-term changes in fear memory and anxiety-like behavior. Psychoneuroendocrinology 104, 300–307. doi:10.1016/j.psyneuen.2019.03.016.

Lalanza, J. F., Caimari, A., del Bas, J. M., Torregrosa, D., Cigarroa, I., Pallàs, M., et al. (2014). Effects of a post-weaning cafeteria diet in young rats: metabolic syndrome, reduced activity and low anxiety-like behaviour. PLoS ONE 9, e85049. doi:10.1371/journal.pone.0085049.

Leigh, S.-J., Kaakoush, N. O., Bertoldo, M. J., Westbrook, R. F., and Morris, M. J. (2020). Intermittent cafeteria diet identifies fecal microbiome changes as a predictor of spatial recognition memory impairment in female rats. Translational Psychiatry 10, 1–12. doi:10.1038/s41398-020-0734-9.

Li, J.-M., Yu, R., Zhang, L.-P., Wen, S.-Y., Wang, S.-J., Zhang, X.-Y., et al. (2019). Dietary fructose-induced gut dysbiosis promotes mouse hippocampal neuroinflammation: a benefit of short-chain fatty acids. Microbiome 7, 98. doi:10.1186/s40168-019-0713-7.

Mager, D. R., Mazurak, V., Rodriguez-Dimitrescu, C., Vine, D., Jetha, M., Ball, G., et al. (2013). A meal high in saturated fat evokes postprandial dyslipemia, hyperinsulinemia, and altered lipoprotein expression in obese children with and without nonalcoholic fatty liver disease. JPEN J Parenter Enteral Nutr 37, 517–528. doi:10.1177/0148607112467820.

Martínez, M. C., Villar, M. E., Ballarini, F., and Viola, H. (2014). Retroactive interference of object-in-context long-term memory: role of dorsal hippocampus and medial prefrontal cortex. Hippocampus 24, 1482–1492. doi:10.1002/hipo.22328.

McNamara, M. P., Singleton, J. M., Cadney, M. D., Ruegger, P. M., Borneman, J., and Garland, T. (2021). Early-life effects of juvenile Western diet and exercise on adult gut microbiome composition in mice. Journal of Experimental Biology. doi:10.1242/jeb.239699.

McNeilly, A. D., Gao, A., Hill, A. Y., Gomersall, T., Balfour, D. J. K., Sutherland, C., et al. (2016). The effect of dietary intervention on the metabolic and behavioural impairments generated by short term high fat feeding in the rat. Physiology & Behavior 167, 100–109. doi:10.1016/j.physbeh.2016.08.035.

Neri, D., Martinez-Steele, E., Monteiro, C. A., and Levy, R. B. (2019). Consumption of ultraprocessed foods and its association with added sugar content in the diets of US children, NHANES 2009-2014. Pediatr Obes 14, e12563. doi:10.1111/ijpo.12563.

Noble, E. E., Hsu, T. M., Jones, R. B., Fodor, A. A., Goran, M. I., and Kanoski, S. E. (2017a). Early-Life Sugar Consumption Affects the Rat Microbiome Independently of Obesity. J Nutr 147, 20–28. doi:10.3945/jn.116.238816.

Noble, E. E., Hsu, T. M., and Kanoski, S. E. (2017b). Gut to Brain Dysbiosis: Mechanisms Linking Western Diet Consumption, the Microbiome, and Cognitive Impairment. Front Behav Neurosci 11. doi:10.3389/fnbeh.2017.00009.

Noble, E. E., Hsu, T. M., Liang, J., and Kanoski, S. E. (2019). Early-life sugar consumption has long-term negative effects on memory function in male rats. Nutr Neurosci 22, 273–283. doi:10.1080/1028415X.2017.1378851.

Noble, E. E., and Kanoski, S. E. (2016). Early life exposure to obesogenic diets and learning and memory dysfunction. Curr Opin Behav Sci 9, 7–14. doi:10.1016/j.cobeha.2015.11.014.

Noble, E. E., Mavanji, V., Little, M. R., Billington, C. J., Kotz, C. M., and Wang, C. (2014). Exercise reduces diet-induced cognitive decline and increases hippocampal brain-derived neurotrophic factor in CA3 neurons. Neurobiol Learn Mem 114, 40–50. doi:10.1016/j.nlm.2014.04.006.

Noble, E. E., Olson, C. A., Davis, E., Tsan, L., Chen, Y.-W., Schade, R., et al. (2021). Gut microbial taxa elevated by dietary sugar disrupt memory function. Translational Psychiatry.

Pindjakova, J., Sartini, C., Lo Re, O., Rappa, F., Coupe, B., Lelouvier, B., et al. (2017). Gut Dysbiosis and Adaptive Immune Response in Diet-induced Obesity vs. Systemic Inflammation. Front Microbiol 8. doi:10.3389/fmicb.2017.01157.

Provensi, G., Schmidt, S. D., Boehme, M., Bastiaanssen, T. F. S., Rani, B., Costa, A., et al. (2019). Preventing adolescent stress-induced cognitive and microbiome changes by diet. Proc. Natl. Acad. Sci. U.S.A. 116, 9644–9651. doi:10.1073/pnas.1820832116.

Qian, L., Huang, J., and Qin, H. (2020). Probiotics and dietary intervention modulate the colonic mucosa-associated microbiota in high-fat diet populations. Turk J Gastroenterol 31, 295–304. doi:10.5152/tjg.2020.19013.

Reichelt, A. C. (2016). Adolescent Maturational Transitions in the Prefrontal Cortex and Dopamine Signaling as a Risk Factor for the Development of Obesity and High Fat/High Sugar Diet Induced Cognitive Deficits. Front Behav Neurosci 10, 189. doi:10.3389/fnbeh.2016.00189.

Romo-Araiza, A., Gutiérrez-Salmeán, G., Galván, E. J., Hernández-Frausto, M., Herrera-López, G., Romo-Parra, H., et al. (2018). Probiotics and Prebiotics as a Therapeutic Strategy to Improve Memory in a Model of Middle-Aged Rats. Front. Aging Neurosci. 10. doi:10.3389/fnagi.2018.00416.

Safari, Z., Monnoye, M., Abuja, P. M., Mariadassou, M., Kashofer, K., Gérard, P., et al. (2019). Steatosis and gut microbiota dysbiosis induced by high-fat diet are reversed by 1-week chow diet administration. Nutrition Research 71, 72–88. doi:10.1016/j.nutres.2019.09.004.

Saiyasit, N., Chunchai, T., Prus, D., Suparan, K., Pittayapong, P., Apaijai, N., et al. (2020). Gut dysbiosis develops before metabolic disturbance and cognitive decline in high-fat diet– induced obese condition. Nutrition 69, 110576. doi:10.1016/j.nut.2019.110576.

Shang, Y., Khafipour, E., Derakhshani, H., Sarna, L. K., Woo, C. W., Siow, Y. L., et al. (2017). Short Term High Fat Diet Induces Obesity-Enhancing Changes in Mouse Gut Microbiota That are Partially Reversed by Cessation of the High Fat Diet. Lipids 52, 499–511. doi:10.1007/s11745-017-4253-2.

Shepherd, J. K., Grewal, S. S., Fletcher, A., Bill, D. J., and Dourish, C. T. (1994). Behavioural and pharmacological characterisation of the elevated “zero-maze” as an animal model of anxiety. Psychopharmacology (Berl) 116, 56–64. doi:10.1007/BF02244871.

South, T., Westbrook, F., and Morris, M. J. (2012). Neurological and stress related effects of shifting obese rats from a palatable diet to chow and lean rats from chow to a palatable diet. Physiology & Behavior 105, 1052–1057. doi:10.1016/j.physbeh.2011.11.019.

Tsan, L., Décarie-Spain, L., Noble, E. E., and Kanoski, S. E. (2021). Western Diet Consumption During Development: Setting the Stage for Neurocognitive Dysfunction. Front. Neurosci. 15. doi:10.3389/fnins.2021.632312.

Wang, H., Liu, C., Liu, Z., Wang, Y., Ma, L., and Xu, B. (2020). The different dietary sugars modulate the composition of the gut microbiota in honeybee during overwintering. BMC Microbiology 20, 61. doi:10.1186/s12866-020-01726-6.

Wang, J., Shang, L., Light, K., O’Loughlin, J., Paradis, G., and Gray-Donald, K. (2015). Associations between added sugar (solid vs. liquid) intakes, diet quality, and adiposity indicators in Canadian children. Applied Physiology, Nutrition, and Metabolism. doi:10.1139/apnm-2014-0447.

Wang, Y., Guglielmo, D., and Welsh, J. A. (2018). Consumption of sugars, saturated fat, and sodium among US children from infancy through preschool age, NHANES 2009-2014. Am J Clin Nutr 108, 868–877. doi:10.1093/ajcn/nqy168.

Yang, Y., Zhong, Z., Wang, B., Xia, X., Yao, W., Huang, L., et al. (2019). Early-life high-fat diet-induced obesity programs hippocampal development and cognitive functions via regulation of gut commensal Akkermansia muciniphila. Neuropsychopharmacology 44, 2054–2064. doi:10.1038/s41386-019-0437-1.

Zhang, C., Zhang, M., Pang, X., Zhao, Y., Wang, L., and Zhao, L. (2012). Structural resilience of the gut microbiota in adult mice under high-fat dietary perturbations. ISME J 6, 1848–1857. doi:10.1038/ismej.2012.27.

