## Supplementary figures and images for "Early life Western diet-induced memory impairments and gut microbiome changes in female rats are long-lasting despite healthy dietary intervention"

### Supplemental Figure 4

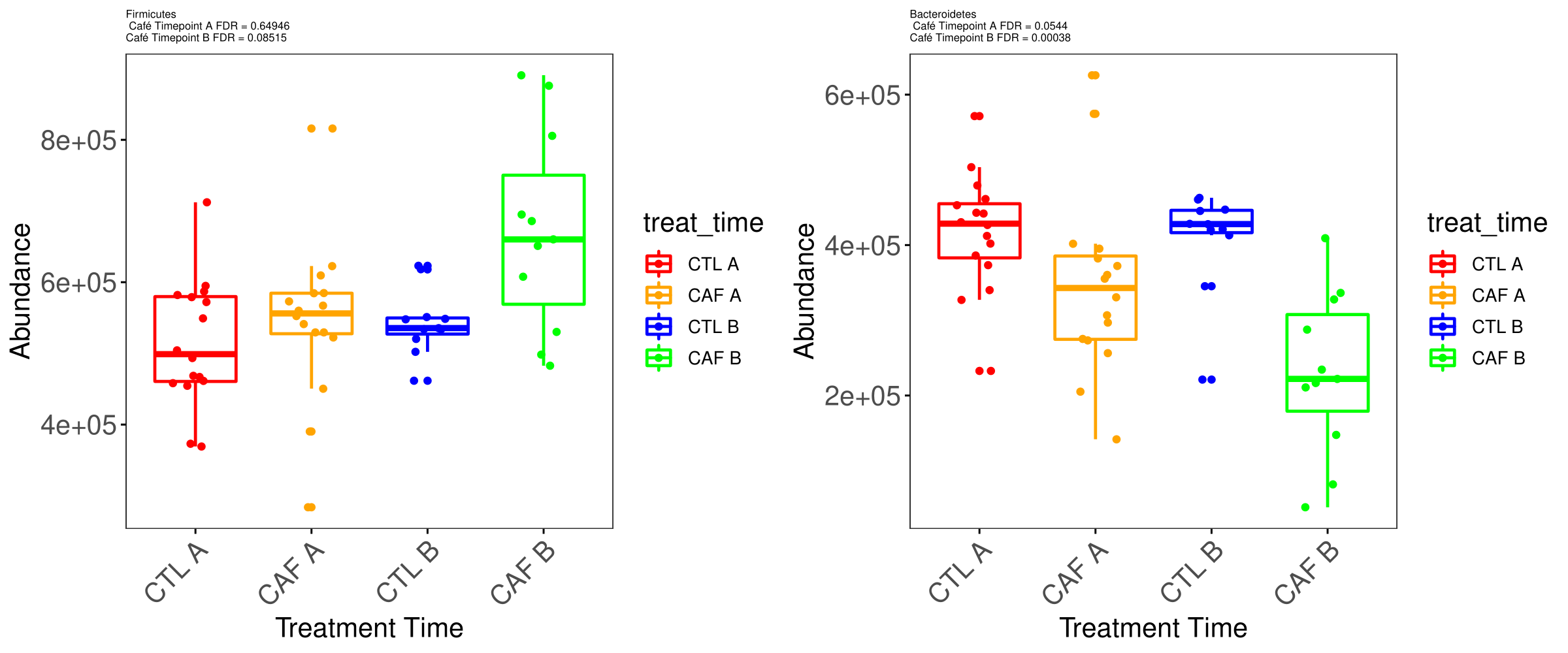

### Supplemental Figure 5

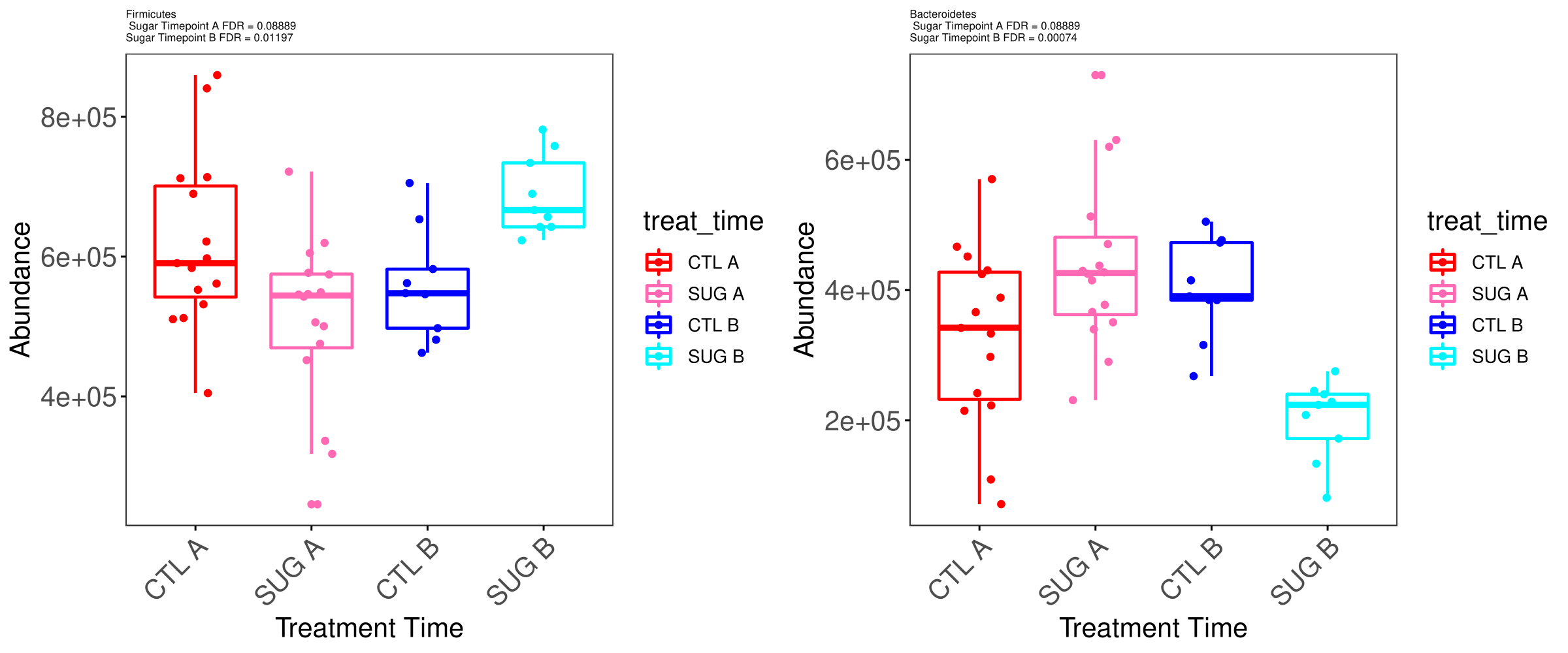

### Supplemental Figure 6

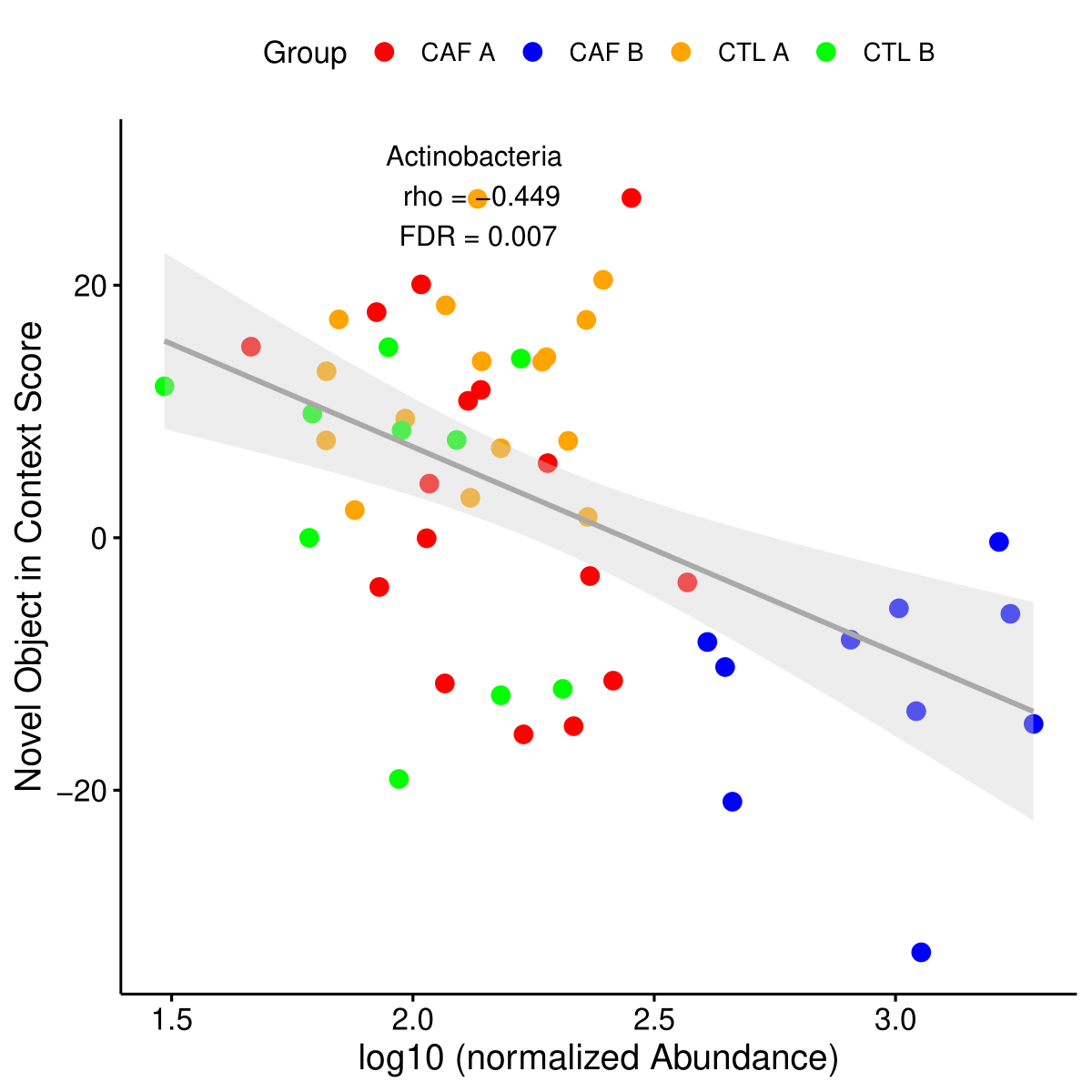
